# Fear Learning–Induced Brain Dynamics Predict Individual Extinction Memory Expression following Transcranial Magnetic Stimulation

**DOI:** 10.64898/2026.03.20.713275

**Authors:** Kai Zhang, Lihan Cui, Isabel Moallem, Hani Meelad, Zakiya Atiyah, Muhammad Badarnee, Isabella Mason, Zhenfu Wen, Mark S. George, Mohammed R. Milad

## Abstract

Fear learning and extinction unfold as time-dependent processes. Herein, we examined how fear learning dynamically reorganizes brain activity immediately after learning, and whether such reorganization can be modulated with TMS application during extinction learning to prospectively predict extinction-memory expression. Eighty-seven healthy adults completed a three-day Pavlovian threat-learning protocol with resting-state fMRI acquired before and after conditioning (Day 1), dorsolateral prefrontal cortex (DLPFC) transcranial magnetic stimulation (TMS) applied during extinction learning (Day 2), and fMRI during extinction recall and renewal (Day 3). Using coactivation pattern analysis with a hidden Markov model within a 24-nodes threat-circuit parcellation, we identified a fear-learning–induced brain state characterized by global threat-circuit coactivation with heightened engagement and transition uncertainty post conditioning, and a progressive increase in engagement across post-conditioning. Critically, conditioning-induced functional connectivity reorganization within this state predicted individual differences in extinction recall- and renewal-related brain activation under TMS-modulated extinction (cross-validated; recall r = 0.47, p = 0.001; renewal r = 0.37, p = 0.01; permutation-tested), but not under natural extinction. Similar associations were observed between neural features and behavioral expression. These findings demonstrate that fear learning reshapes spontaneous brain-state dynamics and that such learning-induced reorganization serves as an interpretable biomarker for neuromodulation-linked extinction-memory expression.

## Introduction

Understanding the underlying neural mechanisms of how conditioned threat memories are generated, consolidated, and updated is central to explaining the neurobiology of human defensive behavior, and to informing interventions for anxiety- and trauma-related disorders(1–3). Despite extensive progress, much of the neuroimaging literature has predominantly emphasized identifying the constituent circuitry engaged by fear-conditioning paradigms, or characterizing learning-related changes using static measures of task-evoked activation or functional connectivity (4–10). However, fear learning and extinction unfold as time-dependent processes, during which cognitive representations and memory traces are continuously reorganized and consolidated. A rich literature from the rodent models implementing fear conditioning and extinction support this view. For example, the engagement of the hippocampus, amygdala, and ventromedial prefrontal cortices- key nodes in fear conditioning and its extinction, is temporally transient during learning (11–18). Consolidation windows after learning have also been shown to occur immediately after learning and gradually fade over time (19). As such, conventional approaches in human neuroimaging research often capture the endpoint of learning-related neural changes, while providing limited access to the moment-to-moment trajectories through which these changes emerge (20–22).

Consistent with a broader shift in systems neuroscience, recent frameworks are increasingly moving beyond static descriptions toward dynamic models that characterize cognition and learning in terms of transitions -through time during and after learning- among a set of recurrent brain states (23–26). For example, Sendi et al demonstrated that alterations in the engagement and stability of specific brain states are associated with subsequent symptom expression following trauma exposure, highlighting the clinical relevance of dynamic state organization in the early aftermath of learning-related stress (27). Longitudinal work from Awada et al further showed that reproducible coactivation patterns constitute stable brain-state signatures linked to motivational and affective disturbances in psychiatric populations (28). Complementarily, clinical trial research from Cai et al revealed that treatment response in major depressive disorder and bipolar disorder is reflected in reorganization of brain-state engagement and transition probabilities (29). These findings illustrate that characterizing state-to-state switching dynamics provides a principled framework for interrogating how cognitive processes and neuromodulation reshape functional brain organization over time. Furthermore, translating basic neuroscience to clinical translational practice is equally important, for example, in describing whether such dynamically defined neural mechanisms can be leveraged as predictive biomarkers to prospectively capture individual variability in treatment response and neuromodulation efficacy (30–32).

Against this background, the present work addresses two important and unresolved questions. First, how fear conditioning dynamically reconfigures the engagement, persistence, and switching structure of spontaneous brain states immediately after learning. Second, whether learning-induced brain-state reorganization predicts subsequent extinction-memory expression, and whether this relationship differs between neuromodulation-enhanced and standard extinction learning procedures. Withing this process, transcranial magnetic stimulation (TMS) as a widely used neuromodulation approach for probing the causal role of prefrontal control circuitry in fear-memory updating. By modulating prefrontal engagement during extinction learning, TMS can selectively influence the modification of threat memories and the retention of extinction, offering an experimental handle on the regulatory processes that govern fear-memory plasticity(33–38). Yet, it remains unresolved whether fear-learning–induced brain-state reorganization can be leveraged as an individual-level neural marker for prospectively predicting extinction-memory expression, and whether the predictive relevance of such reorganization depends on neuromodulation intervention versus natural extinction.

To begin to fill in the gap, we implemented a three-day Pavlovian fear-learning and extinction protocol combined with neuromodulation and data-driven computational modeling (Figure 1). On Day 1, 87 healthy participants completed a Pavlovian fear-conditioning paradigm, in which conditioned threat associations were learned by pairing visual cues of distinct colors (CS+) with aversive electric shocks, while other cues (CS−) were never reinforced. On Day 2, participants underwent extinction learning under transcranial magnetic stimulation (TMS) targeting the dorsolateral prefrontal cortex (DLPFC), enabling experimental modulation of regulatory processes during fear extinction. On Day 3, participants completed memory recall and renewal tasks to probe the expression of conditioned threat memories following extinction and neuromodulation. To characterize the neural dynamics underlying fear learning, we applied coactivation pattern (CAP) analysis to Day 1 resting-state fMRI, identifying a fear-learning–induced brain state that exhibited systematic reorganization of its dynamic properties from pre- to post-conditioning. We then examined whether this learning-induced reorganization predicts subsequent memory expression under TMS navigate extinction and natural extinction on Day 3 using a multivariate predictive framework—regularized canonical correlation analysis (rCCA), and further characterized the neural signatures underlying these brain–brain associations. Moreover, we examined how the fear learning-induced and extinction related brain variates correlate with individual behavioral scores, linking neural dynamics to behavioral measurement across learning process. An illustration of our conceptual and analytic framework is shown in figure 1. Our study not only reveals the dynamic brain reorganization underlying memory consolidation following fear learning, but also demonstrates that these learning-induced dynamics can be formalized as neural biomarkers that effectively predict extinction memory expression after neuromodulation.

**Figure 1.**
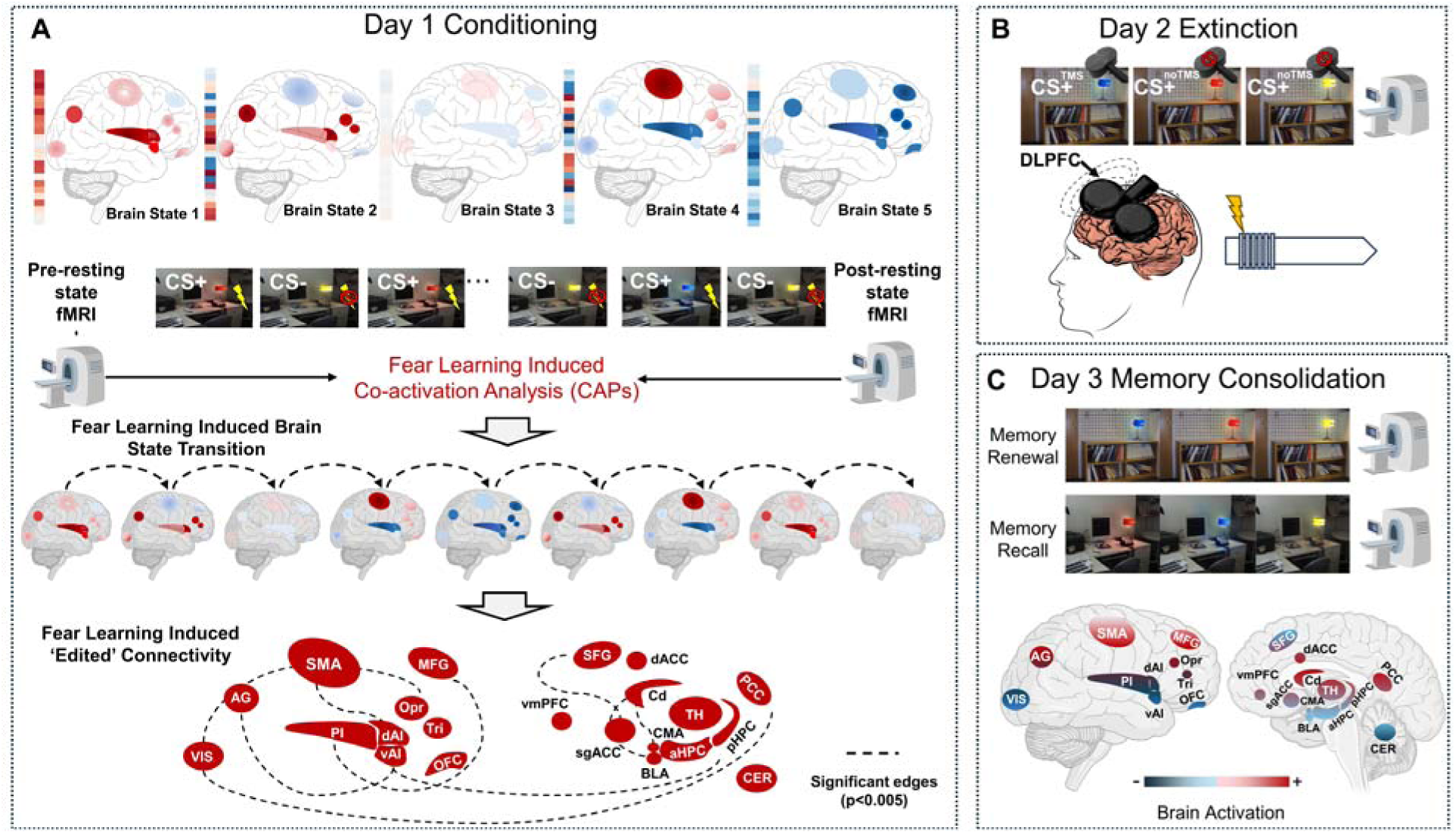
Conceptual and Analytical framework for tracking fear-learning–induced dynamic neural mechanism. Participants completed a three-day experimental protocol (Milad fear learning paradigm) with fear conditioning and extinction. (A). On Day 1, participants underwent a fear conditioning paradigm, with resting-state fMRI acquired before (PRE) and after (POST) conditioning to capture learning-induced changes in spontaneous brain activity. Coactivation pattern (CAP) analysis was applied to PRE-to-POST conditioning resting-state fMRI data to characterize conditioning-induced dynamic brain-state reorganization. (B). On Day 2, participants received TMS-modulated extinction targeting the dorsolateral prefrontal cortex (DLPFC). (C). On Day 3, task for memory recall and fear renewal. Brain activation was used to evaluate extinction memory consolidation. (BLA – basolateral amygdala; CMA – centromedial amygdala; aHPC – anterior hippocampus; pHPC – posterior hippocampus; dAI – dorsal anterior insula; vAI – ventral anterior insula; PI – posterior insular cortex; sgACC – subgenual anterior cingulate cortex; dACC – dorsal anterior cingulate cortex; vmPFC – ventromedial prefrontal cortex; SFG – superior frontal gyrus; MFG – middle frontal gyrus; Tri (IFtri) – triangular part of inferior frontal gyrus; Opr (IFGoper) –opercular part of the inferior frontal gyrus; Precentral – precentral gyrus; Postcentral – Postcentral gyrus; SMA – supplementary motor area; AG – angular gyrus; PCC – posterior cingulate cortex; VIS – visual cortex; TH – thalamus; Cd – caudate nucleus; OFC – Orbital frontal cortex; CER – cerebellum).

## Results

### Fear-learning induced Brain State and its Dynamics Indices

We employed a data-driven approach based on a Hidden Markov Model (HMM) to perform coactivation pattern (CAP) analysis. Resting-state fMRI data acquired before and after fear conditioning, defined within an extended 24-node threat-circuit parcellation(18), were analyzed to characterize the dominant modes of spontaneous brain activity at the population level. Paired t-tests were then used to compare CAP-derived dynamic indicators between pre- and post-conditioning, enabling identification of the dominant brain state modulated by fear learning.

Five distinct recurring patterns of spatial coactivation pattern was defined as basic brain state across all participants. As shown in figure 2A, these states were distinguished by their spatial activation profiles and were labeled based on their predominant circuit engagement. (1) Global threat-circuit activation state, characterized by coordinated positive activation across the majority of regions; (2) Salience-dominant state, marked by strong activation in dAI and angular cortex accompanied by deactivation of vmPFC and sgACC; (3) Silent state, exhibiting minimal activation across regions; (4) Fronto-motor regulatory state, dominated by MFG, SMA, and PCC activation with concurrent suppression of insular regions and (5) Global deactivation state, reflecting widespread attenuation across the circuit.

**Figure 2.**
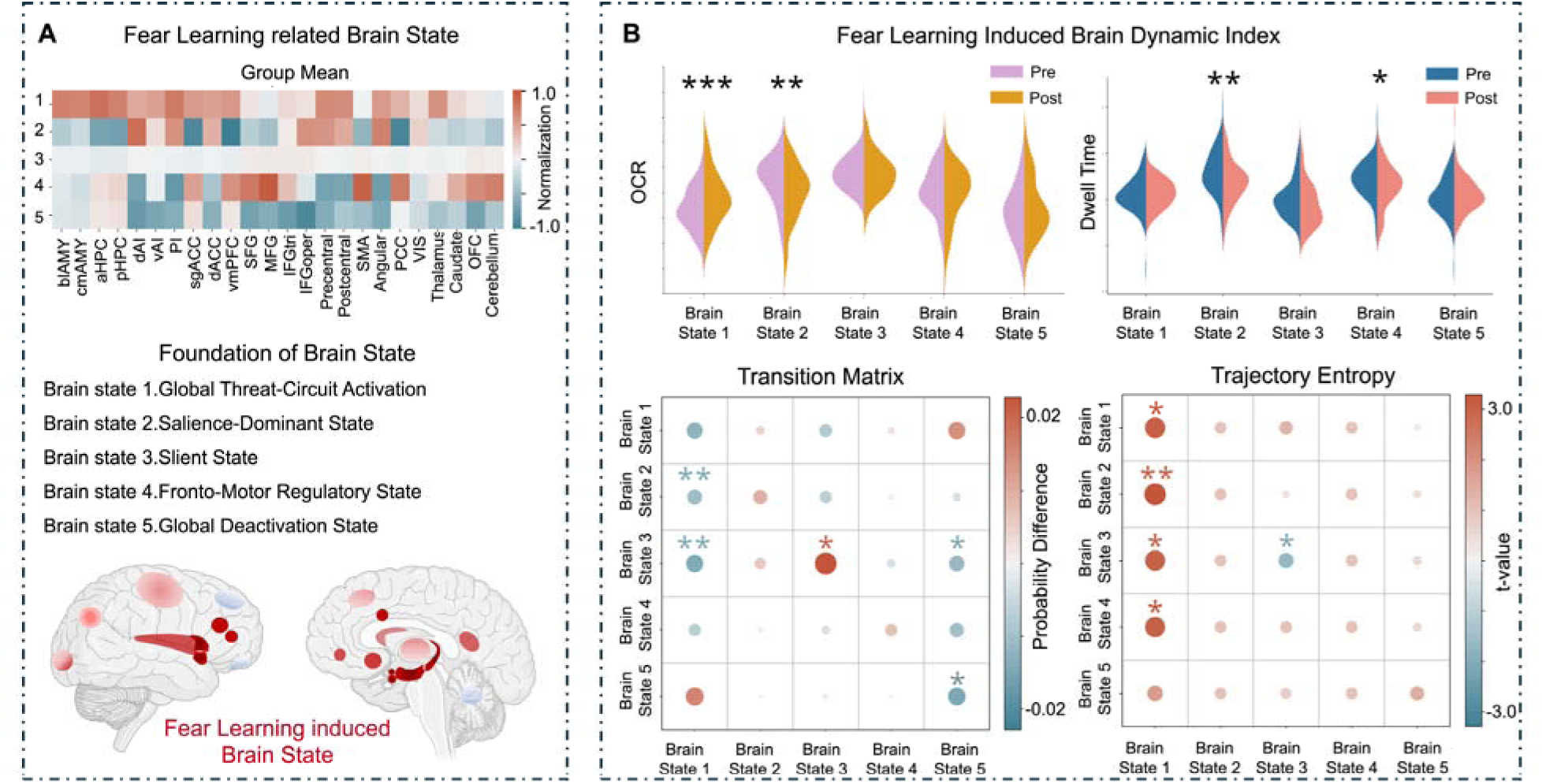
Identification of fear-learning–related dynamic brain states using coactivation pattern (CAP) analysis. (A) Definition of a fear-learning–related brain state based on convergent changes across the four dynamic metrics. The corresponding spatial coactivation pattern is shown within the extended 24-node threat-circuit mask. (B) CAP-derived dynamic metrics characterizing fear-learning–induced brain-state reorganization during Day 1 resting-state fMRI. Four indices were quantified, including occurrence rate (OCR), dwell time, state transition probability matrix, and trajectory entropy, capturing the temporal prevalence, persistence, transition structure, and dynamical complexity of coactivation patterns before and after fear conditioning.

We next examined how fear learning reshaped the temporal dynamics of these states using four complementary dynamic metrics: occurrence rate (OCR) —how often states are engaged, mean dwell time—how long they are sustained (DT), state transition probability—how they switch, and trajectory entropy—how uncertainty or flexibility the switching process is (Figure. 2B). Among the five states, the global threat-circuit activation (State 1) showed a selective and consistent modulation following fear conditioning. Specifically, this state exhibited an increase in occurrence rate (P_FDR_<0.05) from pre-to post-conditioning, indicating more frequent engagement and prolonged persistence after learning. Analysis of the transition probability matrix revealed a biased reorganization of state transitions toward the state 1, with increased transitions from States 2 and 3 into State 1 (P_FDR_<0.05) following conditioning. In parallel, trajectory entropy analyses demonstrated a selective increase in transition entropy involving State 1, reflecting enhanced flexibility and diversification of state-to-state trajectories converging on this fear-related configuration. Notably, entropy increases were observed for transitions from States 1–4 toward State 1 (P_FDR_<0.05), whereas no comparable changes were detected for other states.

### Assessing the temporal dynamics of the Fear Learning-Induced Brain States Formulation Post Fear Conditioning

The change in brain states that are focused on brain circuits to related to fear learning after the acquisition of conditioned fear suggests that this shift might underly the initiation of memory consolidation phase for the recently acquired fear. If so, a temporally-graded shift in these states would be expected. To examine how fear-learning–induced brain dynamics evolve over time following conditioning, we quantified the engagement of the fear-learning–related brain state during the post-conditioning resting period. Specifically, post-conditioning resting-state fMRI time series were segmented into sliding temporal windows of varying lengths. Within each window, we computed the occurrence rate (OCR) of the fear-learning–related brain state, enabling assessment of how state engagement changes as a function of time across multiple temporal resolutions. Across all window lengths, the OCR of the fear-learning–related brain state exhibited a consistent increase over time post the acquisition of conditioned fear conditioning (Figure. 3). Early windows showed relatively low engagement of this state, whereas progressively later windows were characterized by higher OCR, indicating a gradual accumulation of fear-learning–related state engagement over the course of the spontaneous resting period. Mean dwell time (DT) showed a similar increasing trend (Figure S2, Supplementary Materials). This temporal dynamic post-learning supports the idea that this gradual shift might represent the initial phase of memory consolidation associated with threat learning.

**Figure 3.**
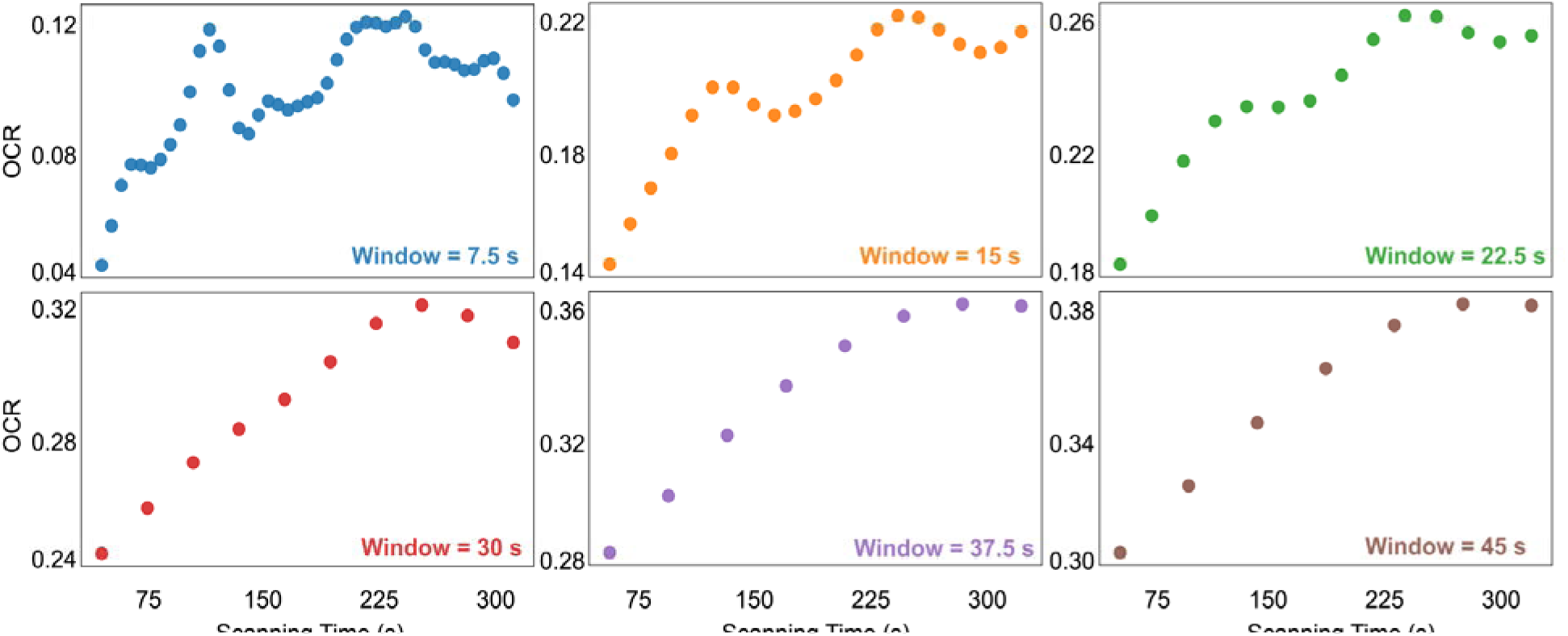
Temporal tracking of fear-learning–induced brain-state engagement during post-conditioning rest. Post-conditioning resting-state fMRI data were segmented using sliding temporal windows of varying lengths. For each window, the occurrence rate of the identified the brain state was computed, indexing its relative engagement over time. The six panels depict OCR trajectories across six window lengths (temporal resolutions), spanning coarse to fine scales. Across temporal resolutions, a progressive increase in the engagement of the fear-learning–related brain state over time during post-conditioning rest.

### Fear-Learning–Induced Brain Reorganization Predicts Extinction Memory

Fear-learning–related connectivity changes (Figure S3, Supplementary Materials) achieved significant out-of-sample predictive relation with the extinction memory-expression variate for both recall and renewal conditions (Figure S4, Supplementary Materials), with r = 0.47, permutation tests p = 0.001 (memory recall) and r = 0.37, permutation tests p = 0.01(memory renewal). Notably, this predictive relationship was specific to the TMS modulation condition. When the same predictive framework was applied to the natural extinction (CS+noTMS), no significant association was observed for either recall or renewal (Supplementary). This dissociation demonstrates that the predictive linkage between fear-learning–induced brain reorganization and later memory-related activation depends on extinction under TMS modulation, rather than reflecting a natural or nonspecific learning effect.

To further characterize the neural signatures underlying these predictive relationships, we examined canonical loadings from the rCCA models to identify features contributing most strongly to each prediction (Figure 5). For the recall condition, connectivity reorganization involved widespread contributions across the extended threat-circuit network. Canonical loading analysis showed that a large proportion of the 24 nodes contributed to the brain-organization variate. On the outcome side, the strongest contributors to the memory-expression variate were localized primarily to the caudate, cerebellum, precentral gyrus, superior frontal gyrus (SFG), and posterior cingulate cortex (PCC). In this model, greater conditioning-induced brain reorganization was associated with reduced activation in these regions, as reflected by negative canonical loadings. In contrast, prediction of renewal-phase activation was driven by a more selective set of learning-related connectivity changes, primarily involving orbitofrontal cortex (OFC), dorsal anterior insula (dAI), ventral anterior insula (vAI), and dorsal anterior cingulate cortex (dACC). On the outcome side, the strongest contributors included IFG opercularis, posterior insula (PI), IFG triangularis, precentral gyrus, and subgenual ACC (sgACC). In this case, greater conditioning-induced brain reorganization was associated with increased activation in these regions, as indicated by positive canonical loadings.

**Figure 4.**
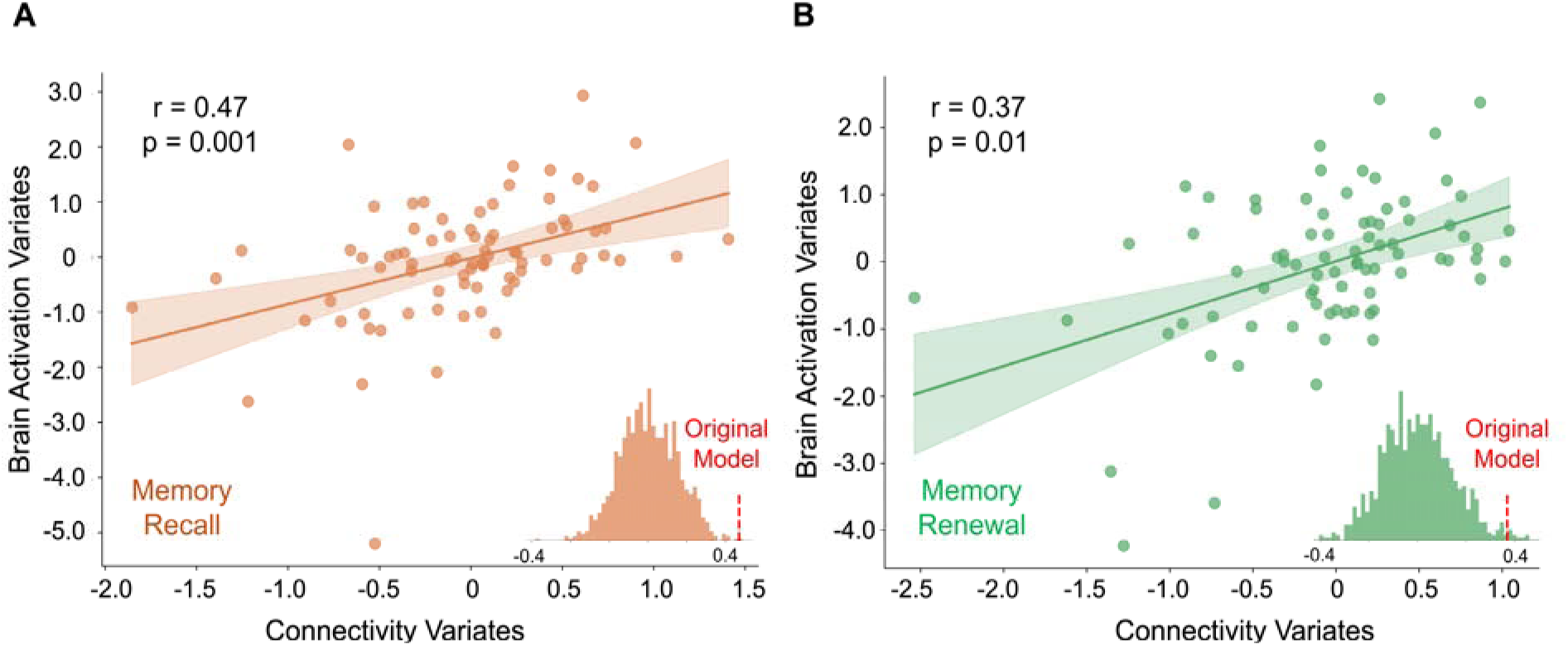
Fear-learning–induced brain reorganization predicts memory recall (A) and renewal (B) brain activation. Connectivity variates was defined by the difference matrix of functional connectivity from post-conditioning minus pre-conditioning brain state 1 for each participant. Brain activation variates is activation value for each participant in recall/renewal phase.

**Figure 5.**
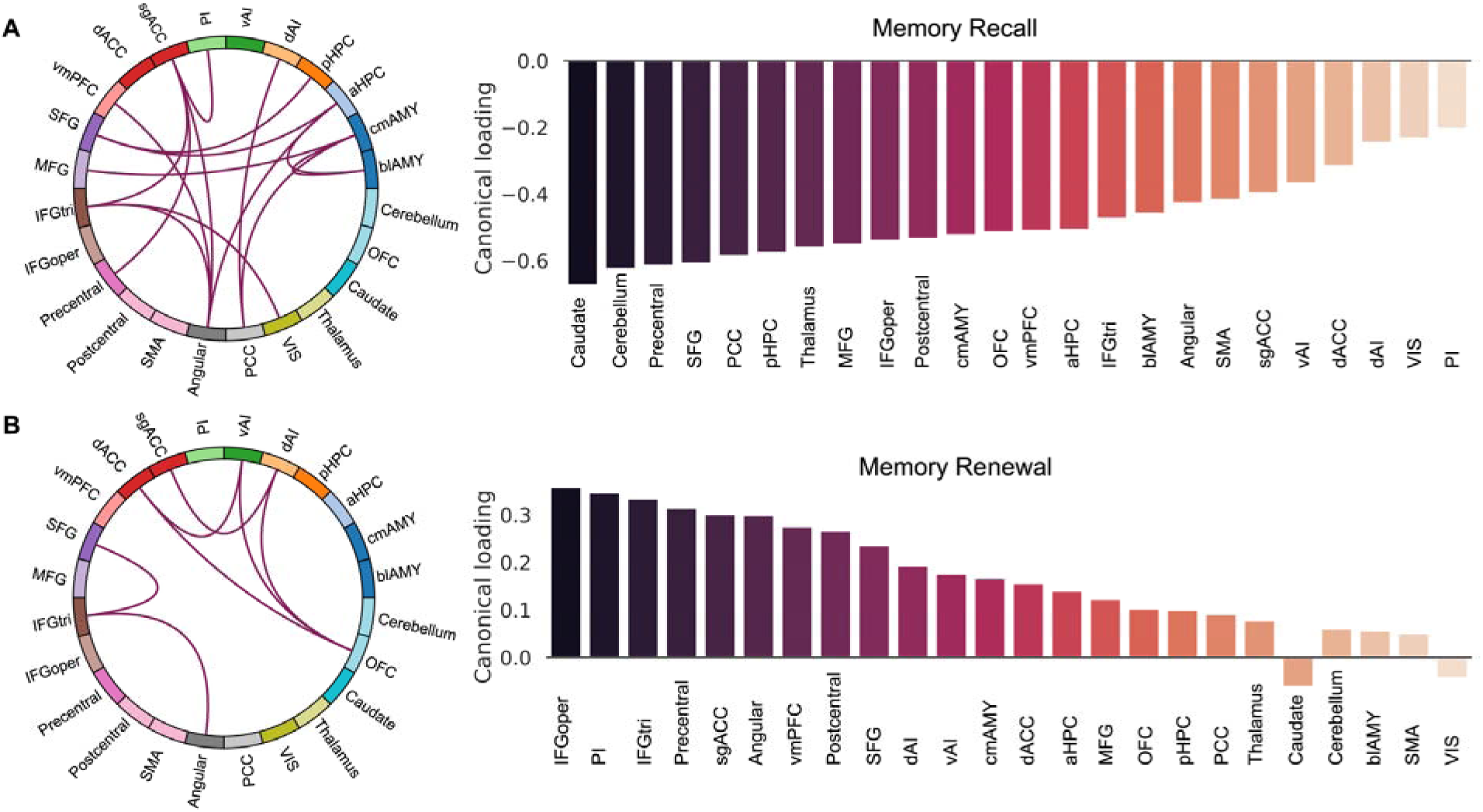
Canonical loading profiles reveal neural signatures linking fear-learning–induced connectivity reorganization to memory recall (A) and renewal (B) brain activation. Canonical loading analysis was performed to characterize the relative contributions of individual features to the canonical variates on both sides of the model.

### Behavioral Results

Behavioral indices of fear reduction were quantified using subjective fear ratings obtained during conditioning and memory expression phase (Figure. 6A–B). For both recall and renewal, CS+ showed a clear reduction of fear level after extinction learning. This reduction was observed in both the TMS and noTMS conditions, whereas fear ratings to the safety cue (CS−) remained stable and close to baseline across all phases. Although the magnitude of fear reduction did not differ significantly between the TMS and noTMS conditions at the group level, the TMS condition exhibited a numerically greater degree of fear attenuation during both groups. This pattern suggests that TMS may enhance the expression of extinction memory, even in the absence of robust group-level behavioral differences.

**Figure 6.**
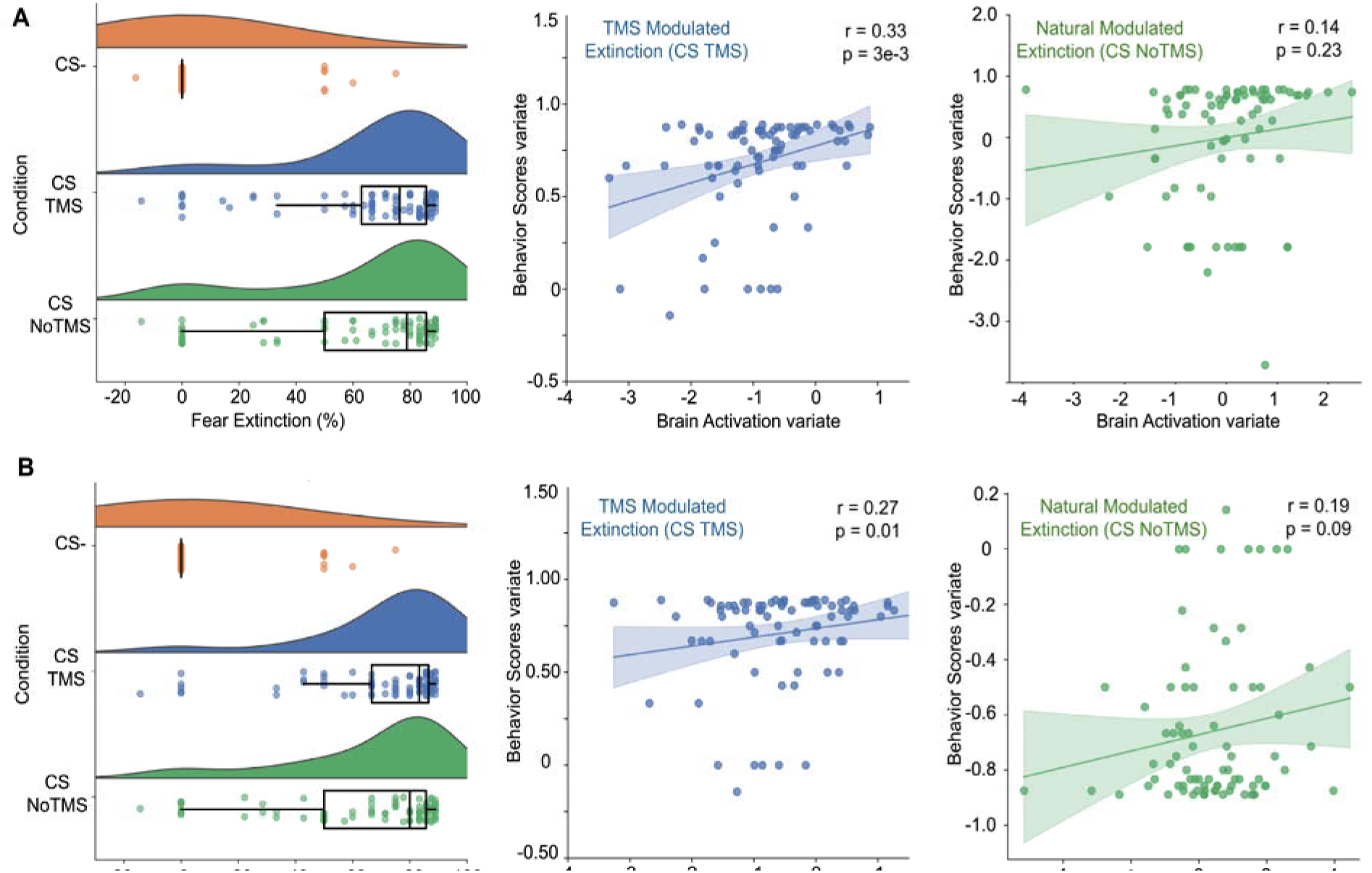
Behavioral expression of extinction memory and its association with neural measures. (A) Subjective fear ratings during the memory recall phase. (B) Subjective fear ratings during the memory renewal phase.

To further examine the relationship between neural dynamics and behavioral expression, we conducted multivariate correlation analyses linking individual differences in behavior to brain-derived measures. As shown in Figure 6A and 6B, TMS modulated extinction memory (recall: r=0.33, p=3e-3; renewal: r=0.27, p=0.01) were significantly associated with subjective fear ratings. In contrast, no significant correlation was observed between natural extinction (CS+NoTMS) memory and behavioral scores (recall: r=0.14, p=0.23; renewal: r=0.19, p=0.09). These findings suggest that although behavioral extinction effects were broadly present across conditions, robust brain–behavior coupling emerged only when extinction was modulated by TMS.

## Discussion

In the present study, we addressed two fundamental questions concerning the neural dynamics of human fear learning and its translational relevance. Specifically, we firstly demonstrated that fear learning selectively reshapes the temporal organization of spontaneous brain states, characterized by increased engagement and altered transition dynamics of a fear-related brain state within a threat-circuits. Secondly, we show that connectivity reorganization induced by fear learning predictively associate with subsequent memory recall and renewal following extinction under TMS-neuromodulation. We establish a dynamic, explainable neural signature of fear learning that not only characterizes how the brain adapts to threat but also serves as a predictive biomarker for extinction memory expression.

Contemporary models of psychopathology increasingly emphasize that maladaptive learning processes in anxiety- and trauma-related disorders arise not solely from abnormalities within static and isolated brain regions, but from dysregulated dynamics across large-scale functional networks(1,39,40). From a cortical dynamics perspective, pathological fear may reflect impaired temporal coordination and flexibility among distributed systems supporting threat detection, salience processing, and regulatory control—manifesting as aberrant engagement, persistence, or switching of large-scale brain states over time(41–46). Despite this theoretical shift, human neuroimaging studies have rarely characterized how fear learning reorganizes brain dynamics at the level of transient brain states, leaving a critical gap in our understanding of the systems-level mechanisms underlying maladaptive fear acquisition and persistence. By leveraging a relatively large sample and a data-driven CAP framework, we demonstrated that fear learning selectively amplifies a distributed threat-related brain state shortly after fear learning. This state was characterized by coordinated engagement of medial temporal, insular, cingulate, and sensorimotor regions, consistent with an integrated network supporting threat encoding, interoceptive salience, and defensive action preparation. Notably, the strongest contributors included the anterior hippocampus, posterior insula, posterior hippocampus, precentral gyrus, and basolateral amygdala that span core components of fear memory formation and expression (Fig S5. Supplement). The hippocampal and amygdala components are central to associative threat learning and contextual memory encoding, the posterior insula supports interoceptive and salience processing of aversive stimuli, and the precentral cortex reflects preparatory motor components of defensive responding. Critically, this fear-learning induced brain state did not merely emerge transiently after conditioning but exhibited a progressive increase in engagement across the post-conditioning resting period. The gradual rise in its engagement over time suggests that fear learning initiates an ongoing consolidation process at the level of large-scale brain dynamics. Rather than reflecting a static imprint of learning, the increasing dominance of this state indicates that spontaneous brain activity continues to reorganize after conditioning, consistent with systems-level consolidation of threat memory.

A central goal of contemporary neuroscience is to link large-scale brain organization to individual variability in treatment response, neuromodulation effects, and cognitive change(30,47–50). Cross-validated predictive modeling has become a powerful framework for capturing the coupling between brain connectivity and functional outcomes at the level of individual subjects (32,51,52). Building on our characterization of fear-learning–induced brain dynamics, we further demonstrate that connectivity reorganization within the fear-learning–related brain state serves as a predictive neural biomarker of subsequent extinction memory retrieval and expression. Specifically, functional connectivity changes from post- to pre-conditioning within this dynamic state significantly predicted individual patterns of brain activation during both memory recall and renewal. Crucially, the predicted activation patterns can be interpreted within a representational framework of threat versus safety processing. For example, by quantifying the relative activation across the 24-node threat-circuit regions, the resulting activation profile can reflect whether the current brain state is dominated by threat-related or safety-related circuitry. Notably, although both CS+TMS and CS+NoTMS conditions exhibited fear extinction, only under TMS did fear-learning–induced brain reorganization predict later recall and renewal activations. The absence of prediction in the CS+NoTMS condition suggests that naturally acquired extinction may engage a convergent, group-level neural implementation. One aspect to note regarding our extinction protocol is that we only administered 4 extinction trials for each of the stimuli (CS+NoTMS and CS+TMS). It is plausible that post-fear learning changes might become associated with natural extinction if additional extinction learning trials occur, a possibility that necessitates testing in future studies. By contrast, TMS modulation appears to promote a more individualized neural expression of extinction memory, such that these activation patterns become systematically predictable from learning-induced network reorganization.

Although both recall and renewal index expressions of extinction memory, they reflect distinct cognitive and contextual processes. Canonical loading analyses revealed dissociable circuit-level signatures for recall and renewal. During memory recall, fear-learning–induced reorganization was distributed across a broad set of connections spanning subcortical, sensorimotor, and medial cortical regions. On the brain-organization variate, prominent contributions arose from connectivity changes involving hippocampal, striatal, cerebellar, and motor-related regions, suggesting that learning-induced reconfiguration within this fear-related state engages distributed systems supporting memory retrieval, action preparation, and contextual integration. Correspondingly, on the memory-expression variate, the most strongly weighted regions included the caudate, cerebellum, precentral gyrus, superior frontal gyrus, and posterior cingulate cortex, indicating that recall-related activation patterns reflect coordinated engagement of motor, cognitive control, and default-mode components. In contrast, renewal exhibited a more selective and regulatory-oriented signature. Fear-learning–related reorganization contributing to renewal prediction was dominated by connectivity changes among orbitofrontal, insular, and cingulate regions, including OFC, dorsal and ventral anterior insula, and dorsal ACC—regions centrally implicated in salience evaluation, interoceptive processing, and regulatory control. On the outcome side, renewal-related activation was most strongly weighted in inferior frontal operculum, posterior insula, inferior frontal gyrus, precentral cortex, and subgenual ACC, highlighting a network configuration consistent with conflict monitoring, affective regulation, and motor readiness during context-driven threat re-expression. The divergence of these neural signatures underscores that extinction memory expression is not monolithic but is instantiated through task- and context-dependent configurations of distributed brain systems shaped by prior fear learning.

Subjective fear ratings indicated that extinction was evident in both stimulation conditions: fear responses decreased from conditioning to the post-extinction memory phase for both CS+TMS and CS+NoTMS, whereas CS− showed minimal change relative to baseline. Although the magnitude of behavioral extinction did not significantly differ between CS+TMS and CS+NoTMS, extinction tended to be stronger in the TMS condition, consistent with a modest facilitative effect of TMS on extinction-memory expression. Importantly, individual differences in behavioral change were systematically related to neural indices derived from the multivariate models. Across recall and renewal, participants showing larger fear-learning–induced reorganization during Day 1 exhibited greater reductions in subjective fear (Figure S6, Supplementary Materials), and stronger extinction-memory–related activation patterns were likewise associated with larger behavioral improvement. In addition, the association between behavioral change and fear-learning–induced reorganization was stronger than that observed for extinction-memory activation, suggesting that the conditioning-phase reconfiguration of functional organization may provide a more proximal neural substrate for inter-individual variability in subsequent extinction outcome.

Several limitations should be acknowledged. First, although our predictive framework demonstrated robust cross-validated performance within the current cohort, external validation in an independent dataset was not possible because no publicly available neuroimaging datasets implement a comparable multi-day fear-conditioning, neuromodulation, and memory-expression paradigm. As a result, the generalizability of the identified dynamic-state biomarkers beyond the present sample remains to be established. Second, the present study focused exclusively on healthy participants. While this approach enabled a controlled characterization of fear-learning–related brain dynamics, it precludes direct inference about clinical populations in whom maladaptive fear learning is most relevant. In particular, it remains unknown whether the fear-learning–induced brain state identified here differs in prevalence, dynamics, or predictive utility between healthy individuals and patients with anxiety- or trauma-related disorders. Future work incorporating patient cohorts will be essential to determine whether alterations in this dynamic brain state represent a mechanistic marker of pathological fear learning and to refine dynamic neural phenotypes associated with impaired adaptive learning.

## Conclusion

In summary, this study provides a dynamic perspective of how human fear learning reshapes large-scale brain organization and how these changes carry predictive value for later memory expression under neuromodulation. By integrating coactivation pattern analysis with multivariate predictive modeling, we identified a fear-learning induced brain state whose engagement and dynamical flexibility increased following conditioning and continued to strengthen over time during post-conditioning. Critically, the magnitude of this learning-induced reorganization predicted individual differences in neural responses during subsequent memory recall and renewal, selectively under TMS-modulated extinction. These findings demonstrate that transient brain dynamics during fear learning are not merely epiphenomenal, but constitute a stable neural biomarker or signature linking learning history to future regulatory outcomes. The present study advance a dynamic neurobiological framework of fear learning and provide a translational bridge from mechanistic brain-state dynamics to individualized prediction of extinction memory expression.

## Method

### Participants

Eighty-seven healthy adults (age range: 18-60 years) was recruited at The University of Texas Health Science Center at Houston (UTHouston). Exclusion criteria included any neurological, psychiatric, or MRI contraindications. All participants provided written informed consent in accordance with the Declaration of Helsinki and protocols approved by the Institutional Review Board of UTHouston.

### Experimental Design

Participants completed a three-day fear-learning and extinction paradigm(14) (Figure. 1, see Supplement for details). On Day 1, participants underwent threat conditioning in which two visual cues (CS+) were paired with an aversive electric shock, whereas a third cue (CS−) was never paired with shock while fMRI data were being acquired. Resting-state fMRI was acquired immediately before and after fear conditioning to capture learning-induced changes in spontaneous brain activity. On Day 2, extinction learning was conducted without shock, during which one CS+ was paired with transcranial magnetic stimulation (TMS) delivered to the left dorsolateral prefrontal cortex (DLPFC), while the other CS+ and the CS− were presented without stimulation. On Day 3, participants completed extinction memory recall and fear renewal phases in which conditioned cues were presented without shock during extinction learning and conditioning contexts, respectively, allowing assessment of extinction memory retrieval and context-dependent fear renewal. On day 3, task-based fMRI data were collected during, extinction recall, fear renewal phases, and resting-state fMRI was acquired pre-task.

### TMS Protocol and Target Selection

Repetitive transcranial magnetic stimulation (rTMS) was administered during extinction learning on Day 2 using a MagPro X100 stimulator equipped with an MC-B70 coil (MagVenture, Farum, Denmark). Stimulation was delivered at a frequency of 20 Hz, with each train lasting 300 ms (six pulses per train) at an intensity of 100% of each participant’s resting motor threshold (rMT). A total of four rTMS trains were delivered across extinction learning, resulting in 24 pulses overall. The brief train duration was selected to model that conducted in rodent studies, and in our prior preliminary human study(1,33). Stimulation was delivered to a target within the left posterior dorsolateral prefrontal cortex (DLPFC), defined as the cortical region showing the strongest and most reliable functional connectivity with the ventromedial prefrontal cortex (vmPFC), a key node implicated in extinction memory. This design constrains stimulation effects to the targeted CS+TMS trials and aims to minimize potential carryover effects across trials. Detailed procedures for target localization, neuronavigation, and rTMS implementation are provided in the Supplementary Methods.

### Neuroimaging Acquisition and Preprocessing

Neuroimaging data were collected on a data were acquired on a 3T Siemens Prisma using a 64-channel head coil. High-resolution T1-weighted anatomical images were acquired for spatial normalization. functional images were collected with a gradient-echo echo-planar imaging (EPI) sequence (TR = 1500 ms; TE = 35 ms; flip angle = 62°; voxel size = 2 × 2 × 2 mm; FOV = 208 mm; 72 slices, interleaved, multi-band acceleration factor = 4).

Data preprocessing was performed using *fMRIPrep* version 20.0.6(53). T1-weighted images underwent intensity nonuniformity correction (N4BiasFieldCorrection, ANTs 2.3.3), skull stripping, tissue segmentation (gray matter, white matter, CSF), and nonlinear normalization to MNI152NLin2009cAsym space (ANTs’ antsRegistration). Functional images were corrected for head motion with mcflirt (FSL), adjusted for slice-timing using 3dTshift (AFNI), and co-registered to the corresponding T1-weighted image using boundary-based registration using flirt (FSL). The resulting functional data were normalized to standard MNI space and resampled to 2 mm isotropic resolution using Lanczos interpolation.

### Dynamic Coactivation Pattern Analysis

We implemented a data-driven coactivation pattern (CAP) analysis that captures momentary configurations of large-scale neural activity(23,54). In this framework, individual BOLD volumes are treated as instantaneous snapshots of whole-brain activation and grouped into a finite set of recurring coactivation states, enabling the identification of transient brain states that emerge during spontaneous activity independent of task timing or experimental condition.

Pre- and post-conditioning resting-state fMRI time series were concatenated across participants, yielding a matrix of size T × R, where T denotes the total number of time points and R the number of brain regions. This concatenated dataset served as the input for CAP modeling, allowing state estimation to be driven by shared spatiotemporal structure across individuals and learning phases.

To determine the appropriate number of coactivation states, clustering solutions ranging from 2 to 14 states were evaluated using complementary model-selection criteria, including the Davies–Bouldin index and the Ray–Turi index, which jointly assess within-state compactness and between-state separability(55). Both metrics converged on an optimal solution comprising five distinct CAP states (Figure S1, Supplementary Materials).

Temporal organization of CAP expression was modeled using a Gaussian hidden Markov model (HMM) with diagonal covariance, trained on the full dataset encompassing both pre- and post-conditioning scans. This probabilistic framework enabled estimation of state-specific temporal trajectories and transitions, thereby capturing learning-related reorganization in dynamic brain-state engagement.

Decoded state sequences were subsequently organized at the subject and phase level and used to derive multiple dynamic indices, including state occurrence rate (OCR), mean dwell time (DT), state transition probabilities (TP), and trajectory entropy (EMT). All these metrics provided a multidimensional characterization of how fear learning reshapes the temporal structure and stability of large-scale brain states(56–58). Detailed mathematical formulations of the HMM, state decoding procedures, and computation of dynamic indices are provided in the Supplementary Materials.

### Estimation of Functional Connectivity Matrix and Brain Activation

Functional connectivity (FC) was estimated using an extended threat-circuit atlas comprising 24 regions of interest(18). For each ROI, regional time series were extracted by averaging the BOLD signal across all voxels within the region. To minimize the impact of head motion, time points with framewise displacement (FD) exceeding 0.3 mm were censored prior to connectivity estimation. For each participant, run-level FC matrices (24 × 24) were computed from resting-state fMRI data using pairwise Pearson correlations between ROI time series. These run-level matrices were then averaged to yield a single subject-level FC matrix for subsequent analyses. The upper triangular elements of each matrix (excluding the diagonal), corresponding to 276 unique functional connections [24 × (24 − 1) / 2], were vectorized and used as input features for predictive modeling.

We used a univariate general linear model (GLM) implemented in the Nistats 0.0.1rc toolbox to estimate activation maps of different stimuli. Given evidence from both human and animal studies shown distinct neural activations across trials even within an experimental phase, we divided trials of each CS type into trial-blocks (4 trials per trial-block) and separately modeled the brain activity to each trial-block. During the conditioning and extinction phases, four trial blocks were modeled for each CS. During the recall and renewal phases, two trial blocks (Early, Late) were modeled for CS+TMS, CS+noTMS, and their corresponding CS− conditions. For each trial-block, the regressor was modeled as boxcar functions time-locked to the CS presentations and convolved with the canonical hemodynamic response function (HRF). Activation estimates from the first trial block (Early) of each condition were used for subsequent predictive analyses.

### Multivariate Prediction of Memory Expression from Fear-Learning–Induced Brain Reorganization

We implemented a multivariate framework based on regularized canonical correlation analysis to test whether fear learning–induced brain reorganization generalizes can predict later memory expression(51,59). rCCA is a multivariate statistical method that identifies pairs of latent components (canonical variates) that maximize the correlation between two high-dimensional variable sets—in this case, state-specific functional connectivity reorganization following fear learning and brain activation patterns during subsequent memory recall or renewal. Compared with classical CCA, rCCA incorporates an L2 regularization term that stabilizes covariance estimation and reduces overfitting, making it well suited for high-dimensional neuroimaging data with limited sample sizes. Model regularization parameters were optimized via a grid-search procedure spanning values from 0 to 1 in increments of 0.05. To prevent information leakage and circular inference, hyperparameter selection was performed exclusively within the training data using a nested 5-fold cross-validation scheme. The optimal regularization parameters identified in the inner loop were then fixed and applied to held-out test data to evaluate out-of-sample predictive performance. Although rCCA yields multiple pairs of canonical variates, primary analyses focused on the first canonical mode, which captures the maximal shared variance between fear-learning–related brain reorganization and later memory-related brain activation. Statistical significance of this canonical association was assessed using a permutation testing procedure, confirming that the observed brain–brain correspondence exceeded chance-level expectations.

### Behavioral Assessment of Fear Extinction

Behavioral indices of fear learning and extinction were assessed using subjective fear ratings collected at key stages of the experimental protocol. Participants were asked to report their perceived level of fear in response to conditioned stimuli using a standardized self-report scale (“Rate your level of fear”), administered following fear conditioning, and again during the post-extinction memory recall and renewal phases.

To quantify individual differences in extinction-related fear reduction, we computed an extinction degree (ED) defined as the normalized change in fear ratings from conditioning to memory expression. Specifically, ED was calculated as the difference between fear ratings obtained after extinction recall phase (i.e., after memory consolidation) and fear ratings obtained immediately after conditioning, divided by the post-conditioning fear level:

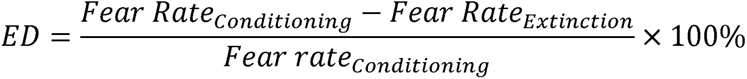

To examine the behavioral relevance of fear-learning–induced brain reorganization, we tested the relationship between ED and the multivariate brain variates derived from the predictive modeling analyses. Specifically, correlations were computed between individual ED scores and the canonical variates capturing fear-learning–related brain reorganization and subsequent memory-related brain activation, allowing assessment of whether neural reorganization during fear learning and its predictive mapping to memory expression covaried with subjective extinction outcomes.

### Statistical Analysis

All dynamic metrics were compared pre versus post conditioning using paired-sample t-tests with FDR-corrected (p < 0.05). In addition, permutation testing (1,000 permutations) was used to assess the statistical significance of predictive associations between fear learning induced brain reorganization and brain activation of extinction memory.

### Code availability statement

The code used for data processing and statistical analyses is publicly available at https://github.com/kaizhang0912/CAP_rCCA_prediction. The repository includes scripts for time-series extraction, functional connectivity computation, co-activation pattern (CAP) analysis (including the calculation of four CAP-related metrics), and rCCA-based prediction analyses. The implementation builds upon publicly available Python libraries including CCA-Zoo https://cca-zoo.readthedocs.io/en/latest/modules/classes.html, Nilearn https://nilearn.github.io/stable/index.html for fMRI data processing, and Seaborn https://seaborn.pydata.org/ for data visualization.

## Data availability statement

The data supporting the findings of this study are available from the corresponding author upon reasonable request. Data sharing is subject to institutional review and must comply with institutional regulations and NIH data-sharing policies to ensure the protection of participant privacy.

---

**Fig S1.**
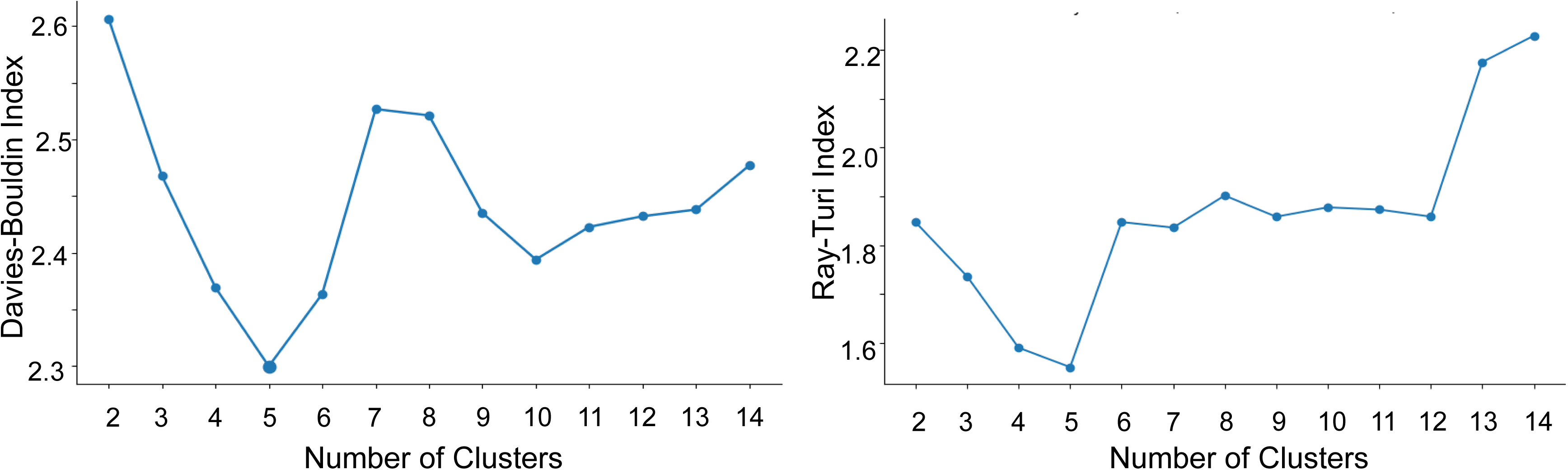

**Fig S2.**
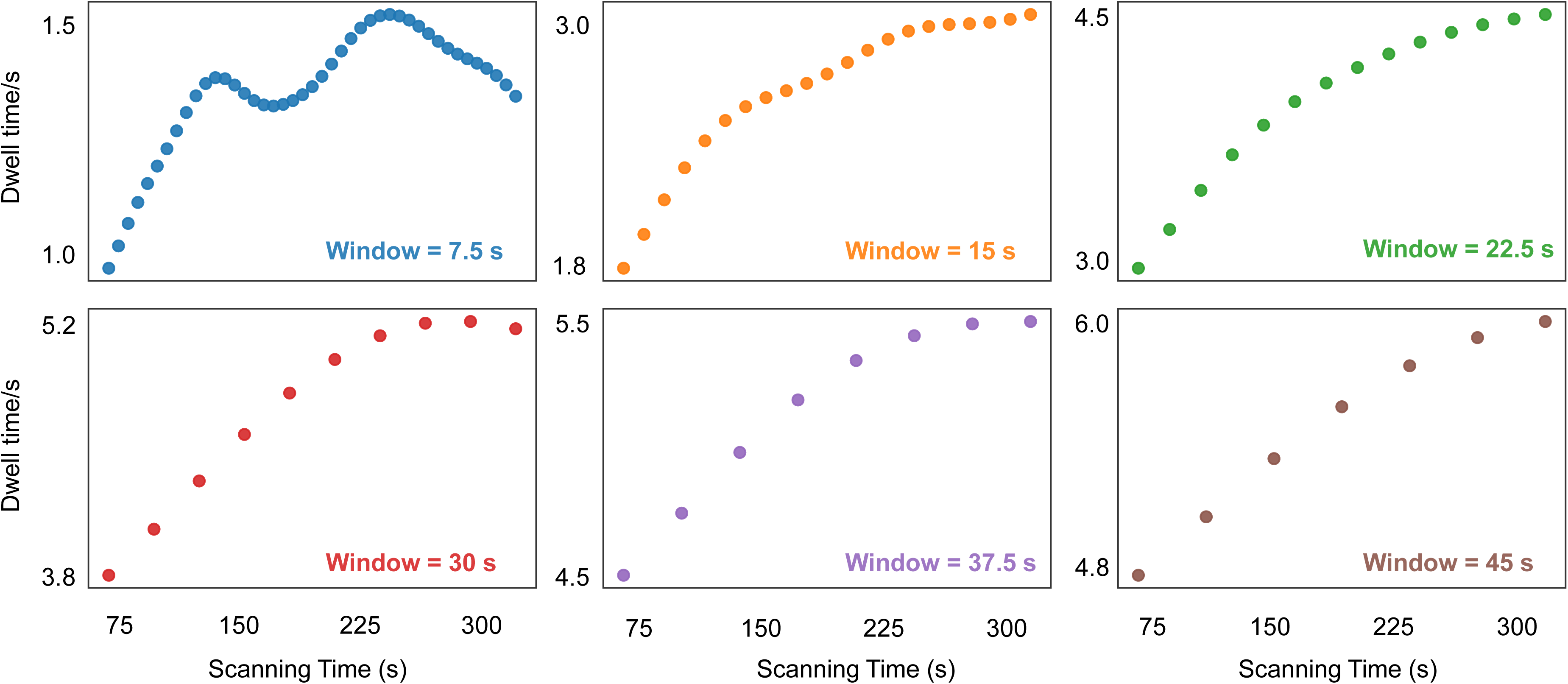

**Fig S3.**
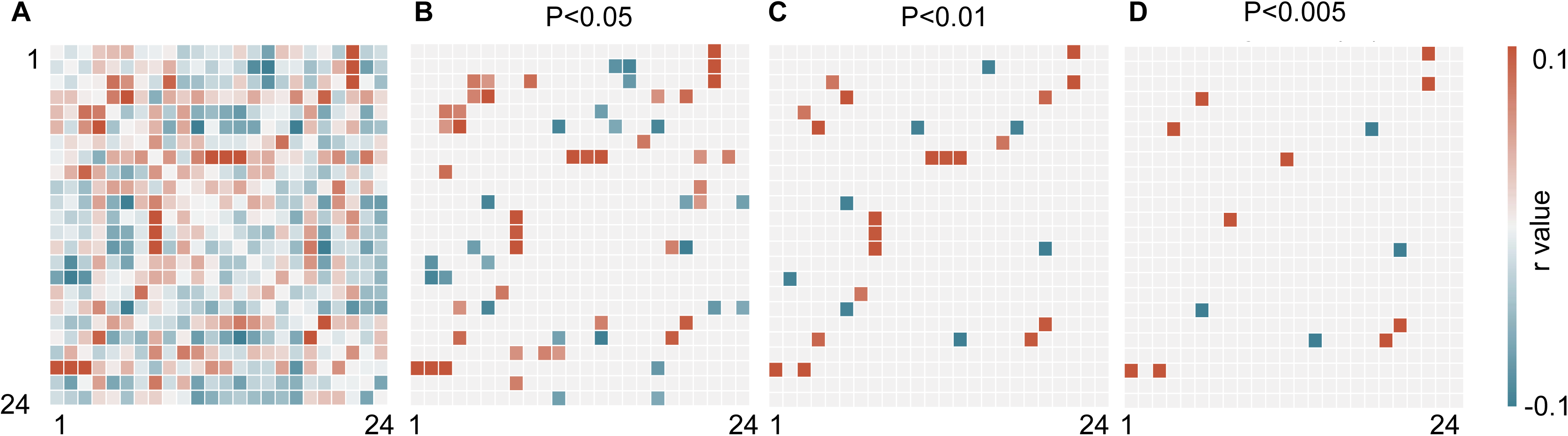

**Fig S4.**
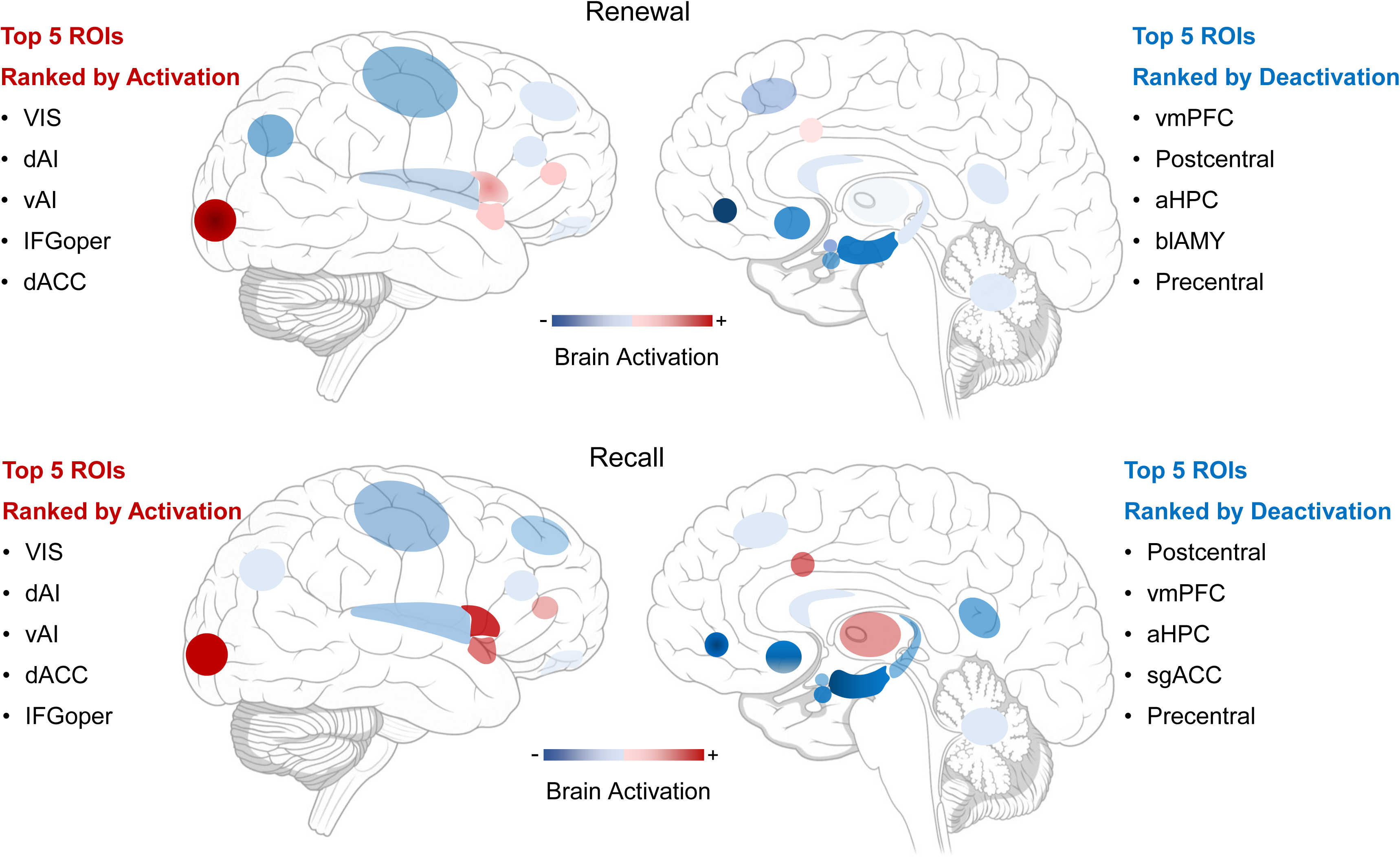

**Fig S5.**
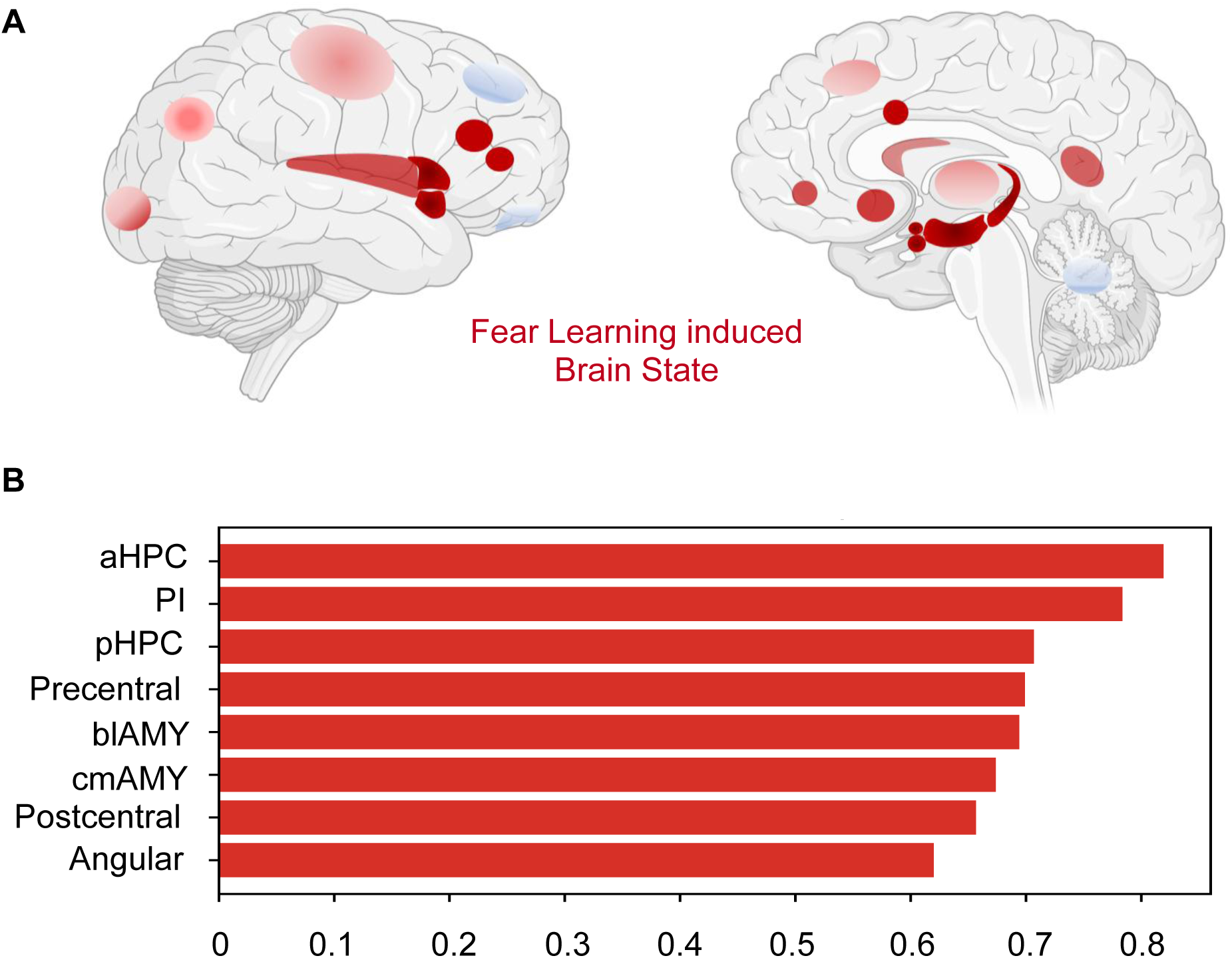

**Fig S6.**
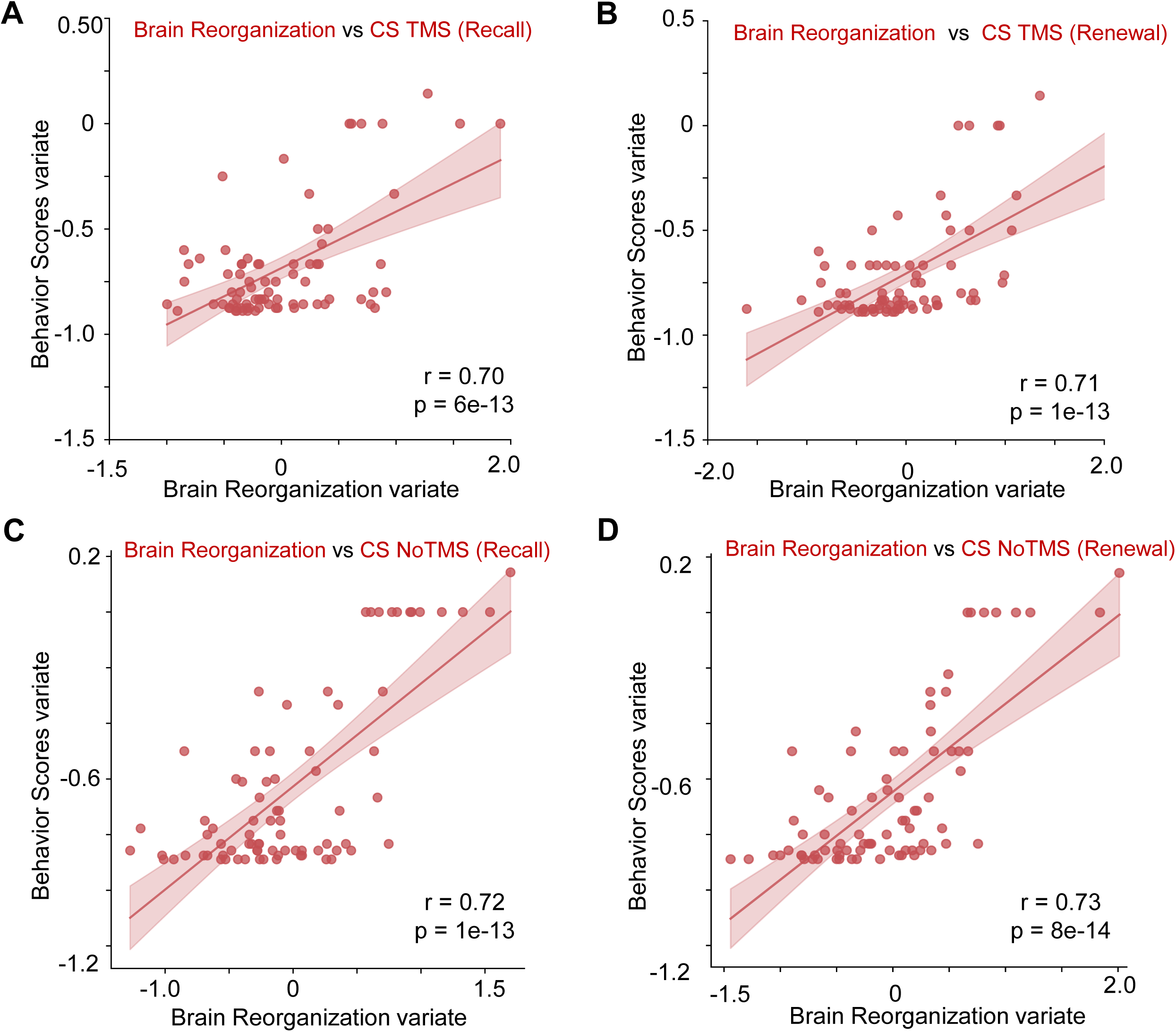

## Reference

1. Milad MR, Quirk GJ (2002): Neurons in medial prefrontal cortex signal memory for fear extinction. Nature 420: 70–74.

2. Milad MR, Quirk GJ (2012): Fear extinction as a model for translational neuroscience: ten years of progress. Annu Rev Psychol 63: 129–151.

3. Olsson A, Phelps EA (2007): Social learning of fear. Nat Neurosci 10: 1095–1102.

4. Greco JA, Liberzon I (2016): Neuroimaging of Fear-Associated Learning. Neuropsychopharmacol 41: 320–334.

5. Fullana MA, Harrison BJ, Soriano-Mas C, Vervliet B, Cardoner N, Àvila-Parcet A, Radua J (2016): Neural signatures of human fear conditioning: an updated and extended meta-analysis of fMRI studies. Mol Psychiatry 21: 500–508.

6. Liu X, Klugah-Brown B, Zhang R, Chen H, Zhang J, Becker B (2022): Pathological fear, anxiety and negative affect exhibit distinct neurostructural signatures: evidence from psychiatric neuroimaging meta-analysis. Transl Psychiatry 12: 405.

7. Klavir O, Prigge M, Sarel A, Paz R, Yizhar O (2017): Manipulating fear associations via optogenetic modulation of amygdala inputs to prefrontal cortex. Nat Neurosci 20: 836–844.

8. Sotres-Bayon F, Sierra-Mercado D, Pardilla-Delgado E, Quirk GJ (2012): Gating of Fear in Prelimbic Cortex by Hippocampal and Amygdala Inputs. Neuron 76: 804–812.

9. Burgos-Robles A, Kimchi EY, Izadmehr EM, Porzenheim MJ, Ramos-Guasp WA, Nieh EH, et al. (2017): Amygdala inputs to prefrontal cortex guide behavior amid conflicting cues of reward and punishment. Nat Neurosci 20: 824–835.

10. Burgos-Robles A, Vidal-Gonzalez I, Quirk GJ (2009): Sustained Conditioned Responses in Prelimbic Prefrontal Neurons Are Correlated with Fear Expression and Extinction Failure. J Neurosci 29: 8474–8482.

11. Phelps EA, LeDoux JE (2005): Contributions of the Amygdala to Emotion Processing: From Animal Models to Human Behavior. Neuron 48: 175–187.

12. Yin S, Liu Y, Petro NM, Keil A, Ding M (2018): Amygdala Adaptation and Temporal Dynamics of the Salience Network in Conditioned Fear: A Single-Trial fMRI Study. eNeuro 5: ENEURO.0445-17.2018.

13. Milad MR, Wright CI, Orr SP, Pitman RK, Quirk GJ, Rauch SL (2007): Recall of Fear Extinction in Humans Activates the Ventromedial Prefrontal Cortex and Hippocampus in Concert. Biological Psychiatry 62: 446–454.

14. Milad MR, Pitman RK, Ellis CB, Gold AL, Shin LM, Lasko NB, et al. (2009): Neurobiological basis of failure to recall extinction memory in posttraumatic stress disorder. Biol Psychiatry 66: 1075–1082.

15. LeDoux J (2003): The emotional brain, fear, and the amygdala. Cell Mol Neurobiol 23: 727–738.

16. Pattwell SS, Duhoux S, Hartley CA, Johnson DC, Jing D, Elliott MD, et al. (2012): Altered fear learning across development in both mouse and human. Proceedings of the National Academy of Sciences 109: 16318–16323.

17. Wen Z, Raio CM, Pace-Schott EF, Lazar SW, LeDoux JE, Phelps EA, Milad MR (2022): Temporally and anatomically specific contributions of the human amygdala to threat and safety learning [no. 26]. Proceedings of the National Academy of Sciences 119: e2204066119.

18. Wen Z, Pace-Schott EF, Lazar SW, Rosén J, Åhs F, Phelps EA, et al. (2024): Distributed neural representations of conditioned threat in the human brain. Nat Commun 15: 2231.

19. Nader K, Schafe GE, Le Doux JE (2000): Fear memories require protein synthesis in the amygdala for reconsolidation after retrieval. Nature 406: 722–726.

20. Garrett DD, Samanez-Larkin GR, MacDonald SWS, Lindenberger U, McIntosh AR, Grady CL (2013): Moment-to-moment brain signal variability: A next frontier in human brain mapping? Neurosci Biobehav Rev 37: 610–624.

21. Mulders PC, van Eijndhoven PF, Schene AH, Beckmann CF, Tendolkar I (2015): Resting-state functional connectivity in major depressive disorder: A review. Neurosci Biobehav Rev 56: 330–344.

22. Sheline YI, Price JL, Yan Z, Mintun MA (2010): Resting-state functional MRI in depression unmasks increased connectivity between networks via the dorsal nexus. Proc Natl Acad Sci U S A 107: 11020–11025.

23. Liu X, Zhang N, Chang C, Duyn JH (2018): Co-activation patterns in resting-state fMRI signals. NeuroImage 180: 485–494.

24. Xie L, Wang X, Lin X, Wen J, Ma T, Leong ATL, Wu EX (2025): Brain-wide resting-state fMRI network dynamics elicited by activation of single thalamic input. Nat Commun 16: 11247.

25. Chen X, Ren H, Tang Z, Zhou K, Zhou L, Zuo Z, et al. (2023): Leading basic modes of spontaneous activity drive individual functional connectivity organization in the resting human brain. Commun Biol 6: 892.

26. Liu Q, Xiong H, Shi W, Di S, Cheng X, Liu N, et al. (2026): Energy inefficiency underpinning brain state dysregulation in individuals with major depressive disorder. Nat Mental Health 1–16.

27. Sendi MSE, Fu Z, Harnett NG, van Rooij SJH, Vergara V, Pizzagalli DA, et al. (2025): Brain dynamics reflecting an intra-network brain state are associated with increased post-traumatic stress symptoms in the early aftermath of trauma. Nat Mental Health 3: 185–198.

28. Awada J, Delavari F, Bolton TAW, Alouf F, Carruzzo F, Kuenzi N, et al. (2026): A Longitudinal and Reproducible Anti-Coactivation Pattern Between the Cerebellum and the Ventral Tegmental Area Is Related to Apathy in Schizophrenia. Biological Psychiatry 99: 124–133.

29. Cai A, Yang J, Guo H, Dong S, Zhao T, Zhou W, et al. (2025): Unraveling spatiotemporal dynamics in transdiagnosis subtypes of major depressive disorder and bipolar disorder: insights from co-activation patterns and treatment response. Mol Psychiatry 1–13.

30. Shen X, Finn ES, Scheinost D, Rosenberg MD, Chun MM, Papademetris X, Constable RT (2017): Using connectome-based predictive modeling to predict individual behavior from brain connectivity. Nat Protoc 12: 506–518.

31. Avissar M, Powell F, Ilieva I, Respino M, Gunning FM, Liston C, Dubin MJ (2017): Functional connectivity of the left DLPFC to striatum predicts treatment response of depression to TMS. Brain Stimulation 10: 919–925.

32. Drysdale AT, Grosenick L, Downar J, Dunlop K, Mansouri F, Meng Y, et al. (2017): Resting-state connectivity biomarkers define neurophysiological subtypes of depression. Nat Med 23: 28–38.

33. Raij T, Nummenmaa A, Marin M-F, Porter D, Furtak S, Setsompop K, Milad MR (2018): Prefrontal Cortex Stimulation Enhances Fear Extinction Memory in Humans. Biological Psychiatry 84: 129–137.

34. Borgomaneri S, Battaglia S, Garofalo S, Tortora F, Avenanti A, Pellegrino G di (2020): State-Dependent TMS over Prefrontal Cortex Disrupts Fear-Memory Reconsolidation and Prevents the Return of Fear. Current Biology 30: 3672–3679.e4.

35. Lynch CJ, Elbau IG, Ng TH, Wolk D, Zhu S, Ayaz A, et al. (2022): Automated optimization of TMS coil placement for personalized functional network engagement. Neuron 110: 3263–3277.e4.

36. Avissar M, Powell F, Ilieva I, Respino M, Gunning FM, Liston C, Dubin MJ (2017): Functional connectivity of the left DLPFC to striatum predicts treatment response of depression to TMS. Brain Stimulation 10: 919–925.

37. McDonald WM, Durkalski V, Ball ER, Holtzheimer PE, Pavlicova M, Lisanby SH, et al. (2011): Improving the antidepressant efficacy of transcranial magnetic stimulation: maximizing the number of stimulations and treatment location in treatment-resistant depression. Depression and Anxiety 28: 973–980.

38. George MS, Wassermann EM, Williams WA, Callahan A, Ketter TA, Basser P, et al. (1995): Daily repetitive transcranial magnetic stimulation (rTMS) improves mood in depression. NeuroReport 6: 1853.

39. Herry C, Johansen JP (2014): Encoding of fear learning and memory in distributed neuronal circuits. Nat Neurosci 17: 1644–1654.

40. LeDoux JE (2000): Emotion Circuits in the Brain. Annual Review of Neuroscience 23: 155–184.

41. Marstaller L, Burianová H, Reutens DC (2016): Dynamic competition between large-scale functional networks differentiates fear conditioning and extinction in humans. NeuroImage 134: 314–319.

42. Uhlhaas PJ, Singer W (2012): Neuronal Dynamics and Neuropsychiatric Disorders: Toward a Translational Paradigm for Dysfunctional Large-Scale Networks. Neuron 75: 963–980.

43. Li X, Zhu D, Jiang X, Jin C, Zhang X, Guo L, et al. (2014): Dynamic functional connectomics signatures for characterization and differentiation of PTSD patients. Hum Brain Mapp 35: 1761–1778.

44. Jin C, Jia H, Lanka P, Rangaprakash D, Li L, Liu T, et al. (2017): Dynamic brain connectivity is a better predictor of PTSD than static connectivity. Human Brain Mapping 38: 4479–4496.

45. Mišić B, Dunkley BT, Sedge PA, Da Costa L, Fatima Z, Berman MG, et al. (2016): Post-Traumatic Stress Constrains the Dynamic Repertoire of Neural Activity. J Neurosci 36: 419–431.

46. Kragel PA, Hariri AR, LaBar KS (2022): The Temporal Dynamics of Spontaneous Emotional Brain States and Their Implications for Mental Health. J Cogn Neurosci 34: 715–728.

47. Wang Z, Goerlich KS, Ai H, Aleman A, Luo Y, Xu P (2021): Connectome-Based Predictive Modeling of Individual Anxiety. Cereb Cortex 31: 3006–3020.

48. Horien C, Floris DL, Greene AS, Noble S, Rolison M, Tejavibulya L, et al. (2022): Functional Connectome–Based Predictive Modeling in Autism. Biological Psychiatry 92: 626–642.

49. Cole MW, Ito T, Bassett DS, Schultz DH (2016): Activity flow over resting-state networks shapes cognitive task activations. Nat Neurosci 19: 1718–1726.

50. Tavor I, Jones OP, Mars RB, Smith SM, Behrens TE, Jbabdi S (2016): Task-free MRI predicts individual differences in brain activity during task performance. Science 352: 216–220.

51. Zhang K, Klumpp H, Jimmy J, Phan KL, Milad MR, Wen Z (2025): Functional Connectivity Predicting Transdiagnostic Treatment Outcomes in Internalizing Psychopathologies. JAMA Netw Open 8: e2530008.

52. Zhao K, Fonzo GA, Xie H, Oathes DJ, Keller CJ, Carlisle NB, et al. (2024): Discriminative functional connectivity signature of cocaine use disorder links to rTMS treatment response. Nat Mental Health 2: 388–400.

53. Esteban O, Markiewicz CJ, Blair RW, Moodie CA, Isik AI, Erramuzpe A, et al. (2019): fMRIPrep: a robust preprocessing pipeline for functional MRI. Nat Methods 16: 111–116.

54. Meer JN van der, Breakspear M, Chang LJ, Sonkusare S, Cocchi L (2020): Movie viewing elicits rich and reliable brain state dynamics. Nat Commun 11: 5004.

55. Vergara VM, Salman M, Abrol A, Espinoza FA, Calhoun VD (2020): Determining the number of states in dynamic functional connectivity using cluster validity indexes. Journal of Neuroscience Methods 337: 108651.

56. An Z, Tang K, Xie Y, Tong C, Liu J, Tao Q, Feng Y (2024): Aberrant resting-state co-activation network dynamics in major depressive disorder. Transl Psychiatry 14: 1–12.

57. Peng X, Liu Q, Hubbard CS, Wang D, Zhu W, Fox MD, Liu H (2023): Robust dynamic brain coactivation states estimated in individuals. Science Advances 9: eabq8566.

58. Chen JE, Chang C, Greicius MD, Glover GH (2015): Introducing co-activation pattern metrics to quantify spontaneous brain network dynamics. NeuroImage 111: 476–488.

59. Mihalik A, Chapman J, Adams RA, Winter NR, Ferreira FS, Shawe-Taylor J, Mourão-Miranda J (2022): Canonical Correlation Analysis and Partial Least Squares for Identifying Brain–Behavior Associations: A Tutorial and a Comparative Study. Biological Psychiatry: Cognitive Neuroscience and Neuroimaging 7: 1055–1067.

