## Supplemental Data 1 for "Fear Learning–Induced Brain Dynamics Predict Individual Extinction Memory Expression following Transcranial Magnetic Stimulation"

### Method

#### Experimental Design

Participants completed a three-day Pavlovian fear-learning and extinction protocol adapted from our previously established paradigm (see Figure 1). Conditioned threat learning was induced using visual cues paired with aversive electric foot shocks (BIOPAC 160 Systems, Goleta, CA).

On Day 1 (fear conditioning), participants were presented with visual stimuli of three different colors. Two colors served as conditioned threat cues (CS+), each delivered in separate blocks, whereas a third color served as a safety cue (CS−) and was presented across both blocks. Each trial began with the presentation of a context image (office) for 3 s, followed by the CS for 6 s. In CS+ trials, a 500 ms shock was delivered at CS offset. CS− trials were never reinforced. The mean inter-trial interval (ITI) was 15 s. Shock intensity was individually calibrated to be highly aversive but not painful.

On Day 2 (extinction learning), participants underwent extinction training in the absence of shock. The same three visual cues were presented across two blocks, with four trials per CS+ and four trials for CS−. During extinction, one CS+ was paired with brief trains of repetitive transcranial magnetic stimulation (rTMS; CS+TMS), whereas the other CS+ and the CS− were not paired with stimulation (CS+NoTMS and CS−). rTMS trains were delivered to the left prefrontal cortex. This within-subject design enabled direct comparison of extinction learning under neuromodulation versus a control condition.

On Day 3, participants completed memory recall and renewal testing. The same conditioned cues were presented without shock in two blocks. Recall trials were conducted in the same contextual environment as extinction learning on Day 2, whereas renewal trials were conducted in the original conditioning context from Day 1, allowing dissociation of extinction memory retrieval from context-dependent threat renewal. The order of CS+TMS and CS+NoTMS blocks, as well as cue–color assignments, was randomized across participants.

### TMS Protocol

#### 1.TMS Administration and Neuronavigation

Transcranial magnetic stimulation (TMS) was administered during extinction learning on Day 2 using a MagPro X100 stimulator equipped with an MC-B70 figure-of-eight coil (MagVenture, Farum, Denmark). Neuronavigation was performed with Brainsight 2.4 (Rogue Research Inc.) to ensure accurate and reproducible coil positioning relative to each participant’s individual brain anatomy throughout the stimulation session.

#### 2.Structural and Functional Image Integration

High-resolution T1-weighted anatomical images acquired during the MRI session were imported into Brainsight for each participant. To guide stimulation target selection, individual anatomical images were integrated with functional connectivity maps derived from task-related fMRI analyses conducted during fear conditioning. Specifically, seed-based functional connectivity maps identifying cortical regions functionally coupled with ventromedial prefrontal cortex (vmPFC) were overlaid onto each participant's anatomical reconstruction.

Skin and full-brain curvilinear surface reconstructions were generated within Brainsight to enable precise visualization of cortical landmarks and to facilitate accurate coil placement and orientation relative to the underlying cortical target.

**3.Connectivity-Informed Target Definition**

Because vmPFC is a deep cortical structure not directly accessible to TMS, a connectivity-informed targeting strategy was employed to identify a surface-accessible cortical site capable of indirectly modulating vmPFC circuitry. Seed-based functional connectivity analyses were performed using fMRI data acquired during fear conditioning. A vmPFC region of interest was defined a priori, and for each participant, the vmPFC time series was correlated with voxel-wise BOLD signals across the brain to generate individual whole-brain functional connectivity maps.

Based on these maps, a target within the left posterior dorsolateral prefrontal cortex (DLPFC) was identified as the cortical region showing the strongest and most reliable functional connectivity with vmPFC. This region was selected as the primary stimulation target with the rationale that stimulation at this site would induce secondary modulation of vmPFC through established functional network pathways relevant to fear regulation.

**4.Landmark Registration and Target Localization**

Standard cranial landmarks—including the left and right preauricular points, nasion, and nose tip—were identified for each participant and registered to establish accurate coregistration between head position and MRI space. Registration accuracy was verified using Brainsight's built-in validation procedure prior to stimulation.

Two stimulation sites were defined for each participant:

- Motor threshold (MT) site, localized over the hand knob region of the primary motor cortex and used to determine the individual resting motor threshold.
- DLPFC stimulation site, defined using the connectivity-informed procedure described above and targeted to induce secondary modulation of vmPFC circuitry during extinction learning.

**5.Stimulation Parameters**

The TMS coil was calibrated using Brainsight's tool calibration procedure prior to each session. Resting motor threshold (rMT) was determined by stimulating the motor cortex and identifying the minimum intensity that elicited visible hand muscle twitches in at least 3 out of 10 trials. Stimulation intensity for the extinction protocol was set to 100% of the individual rMT.

During extinction learning, repetitive TMS (rTMS) was delivered at:

- Frequency: 20 Hz
- Train duration: 300 ms (6 pulses per train)
- Intensity: 100% rMT
- Total number of trains: 4 (24 pulses total)

The short train duration was selected to confine neuromodulatory effects to the target trial and minimize carryover effects across trials or blocks.

A within-subject control condition was implemented by pairing rTMS with only one conditioned stimulus (CS+TMS), while the other conditioned stimulus (CS+NoTMS) and the safety cue (CS−) were presented without stimulation. Coil positioning was continuously monitored using Brainsight's real-time feedback (bullseye coil-centric view), and stimulation was administered only when spatial error was below 1 mm.

Participants were closely monitored throughout stimulation to ensure comfort and safety. Stimulation was immediately discontinued if participants reported discomfort or intolerance. All procedures adhered to established safety guidelines for TMS application.

**Dynamic Coactivation Pattern Analysis**

To model temporal dynamics, we trained a Gaussian hidden Markov model (HMM) on the entire dataset, which included both pre- and post-conditioning scans from all participants. Each time point was assigned to the latent state with the highest posterior probability, yielding subject-specific state sequences for all participants across all tasks and sessions.

State sequences were then organized into structured dictionaries indexed by subject, task, timepoint, and diagnostic group, and used to derive dynamic metrics including occurrence rate (OCR), mean dwell time (DT), state transition probability matrix (TP), and entropy of Markov trajectories (EMT). Specifically, these indices capture distinct but related aspects of brain dynamics—how often states are engaged (OCR), how long they are sustained (DT), how they switch (TP), and how predictable or uncertain the switching process is (EMT).

**Occurrence.** For each participant, we quantified state occurrence as the proportion of time points assigned to each latent state. Specifically, decoded state sequences were concatenated across tasks and the number of samples in each state was divided by the participant's total number of valid samples after preprocessing, yielding a *k*-dimensional vector of subject-specific occupancy rates that sum to 1.

**Dwell time**. For each participant and task, we computed dwell time from the decoded state sequence by identifying runs of consecutive time points assigned to the same latent state. A run's DT equals its run-length in TRs (i.e., the number of consecutive samples until the state changes), including self-runs of length 1. For each participant and state, we obtain a subject-specific mean DT.

**Transition probabilities**. We estimated transition probabilities empirically from each participant's decoded state. For each participant, we counted all observed transitions between successive time points across tasks (including self-transitions) to form $k \times k$ count matrix. Counts were then row-normalized so that outgoing probabilities from each "from-state" summed to 1, yielding a participant-specific transition probability (TP) matrix.

**Entropy of Markov trajectories.** For each participant, we derived a $k \times k$ times transition probability matrix *P* from empirical counts of adjacent states in the decoded sequence. We first computed the local (row) entropy in bits for each "from-state", $h_i = -\sum_j P_{ij} log P_{ij}$. We then obtained the stationary distribution *μ* (left eigenvector for eigenvalue 1, normalized to sum to 1) and calculated the trajectory entropy matrix *H*, which quantifies the expected cumulative uncertainty along Markov trajectories from state *i* to *j*. This was implemented following standard formulations using the fundamental matrix$(I - P + A)^{-1}$, where A tiles *μ* by rows, with corrections based on the local entropy vector and the chain's entropy rate. For group-level analysis, we summarized EMT at two levels: (i) a scalar EMT per participant obtained by averaging all  entries, and (ii) the full EMT matrix *H* for element-wise comparisons. Higher EMT indicates greater uncertainty/dispersion of the state-transition process. Group-level trajectory entropy differences were visualized using matrix-based bubble plots. For each state pair (i,j), the color of each cell represented the paired t-statistic comparing pre- and post-condition EMT values across participants, while circle size reflected the absolute magnitude of the statistic. Diverging colormaps centered at zero were used to encode directionality of change, with warm colors indicating increases and cool colors indicating decreases. To ensure robust visualization and avoid scaling distortions caused by extreme values, color limits were symmetrically defined based on percentile thresholds of the statistic distribution.

Finally, we computed fear learning induced changes in dynamic features (OCR, DT, TP, EMT) between pre- and post-conditioning sessions.

**Predictive Modeling Using Regularized Canonical Correlation Analysis (rCCA)**

We examined whether fear-learning–induced brain reorganization can predict subsequent extinction-related memory expression, as indexed by task-evoked brain activation and behavioral measures during recall and renewal. To this end, we applied regularized canonical correlation analysis (rCCA) with a nested cross-validation framework to assess multivariate associations between learning-related changes in dynamic brain organization and later neural and behavioral outcomes.

**1. Brain and Outcome Variate Definition**

Two multivariate domains were defined for the rCCA analyses.

Brain organization variate (X).

Based on the coactivation pattern (CAP) analysis, we identified a fear-learning–related brain state that showed systematic reorganization from pre- to post-conditioning. For each participant, we computed state-specific functional connectivity (FC) matrices for this fear-related CAP using resting-state fMRI acquired before and after conditioning. Regional time series were extracted from a predefined 24-node extended threat-circuit parcellation, and pairwise Pearson correlations were used to estimate FC within the CAP-assigned frames.

For each participant, FC matrices were computed separately for the pre- and post-conditioning resting-state scans and then differenced (POST − PRE) to index fear-learning–induced connectivity reorganization. The upper triangular elements of each FC matrix (excluding the diagonal), yielding 276 unique connections, were vectorized and used as the brain organization feature set (X).

Memory-expression variate (Y).

Outcome variables were defined as task-evoked neural activation patterns during Day 3 memory recall and renewal. Activation estimates were derived from task-based fMRI using a general linear model and summarized within the same 24-node threat-circuit atlas. Separate rCCA models were constructed for recall and renewal conditions. In complementary analyses, subjective fear ratings indexing extinction memory expression were also included as behavioral outcome variables.

**2. rCCA Model Specification**

We fitted the rCCA model to identify linear combinations of the brain organization variate (X) and memory-expression variate (Y) that maximized their correlation. rCCA was implemented using the CCA-Zoo Python library (version 2.6.0). Compared to standard

CCA, rCCA incorporates an L2 regularization term on the covariance matrices of both datasets, enhancing numerical stability and reducing overfitting in high-dimensional feature spaces where the number of predictors exceeds the number of observations.

The regularization parameter was optimized within an inner cross-validation loop, with candidate values ranging from 0 to 1 in increments of 0.05.

**3. Nested Cross-Validation Procedure**

To evaluate generalization performance and avoid circularity, we implemented a nested cross-validation framework. In the outer loop, we conducted 10-fold cross-validation repeated five times. In each iteration, nine folds were used for model training and one fold for testing. Canonical weights derived from the training data were applied to the held-out test data to generate predicted canonical variate scores.

Within each outer training set, an inner 5-fold cross-validation procedure was used to select the optimal regularization parameter. Specifically, the training data were split into five inner folds, and for each candidate regularization value, the model was trained on four folds and evaluated on the remaining fold. This process was repeated across all inner folds, and the parameter yielding the highest average performance was selected. The model was then refit using the full outer training set and applied to the outer test set.

This nested design ensured that hyperparameter optimization was strictly confined to the training data, providing an unbiased estimate of out-of-sample predictive performance.

**4. Permutation Testing**

Statistical significance of the predictive associations was assessed using permutation testing. Specifically, we performed 1,000 permutations in which the rows of the outcome matrix (Y) were randomly shuffled prior to cross-validation, while preserving the structure of the brain organization matrix (X). The observed mean out-of-sample canonical correlation was compared against the resulting null distribution to obtain permutation-based p-values.

**5. Canonical Loadings and Neural Signature Analysis**

To characterize the neural features contributing most strongly to the predictive associations, we conducted a canonical loadings analysis using cross-validated predictions. In each fold of the outer cross-validation, canonical variate scores were generated for participants in the test set. These predicted scores were then concatenated across folds to obtain a full set of out-of-sample canonical variate estimates.

Connectivity loadings were computed as the Pearson correlation between the predicted brain-organization variate scores and each functional connectivity feature across participants. Similarly, activation loadings were computed as the Pearson correlation between predicted memory-expression variate scores and regional activation features. Features exceeding a significance threshold of $p < 0.01$ were visualized to delineate the neural signature linking fear-learning–induced brain reorganization to later memory expression.

### Result

#### Numbers of Coactivation States

To select the optimal number of recurrent CAPs, we computed clustering solutions for k=2 through k=14 and evaluated each using two internal validity metrics: the Davies–Bouldin (DB) index and the Ray–Turi (RT) index. In Supplementary Figure S1 (left panel), the DB index measures the ratio of intra-cluster scatter to inter-cluster separation—

reaches its minimum value at k=5. A lower DB score indicates tighter, more distinct clusters, so this nadir suggests that a five-cluster solution most effectively balances within-state cohesion against between-state separation. The RT index (right panel) similarly attains one of its lowest values at k=5, indicating optimal cluster dispersion relative to each centroid. On the strength of this convergence, we fixed the number of CAPs to 5 for all subsequent HMM-CAP analyses.

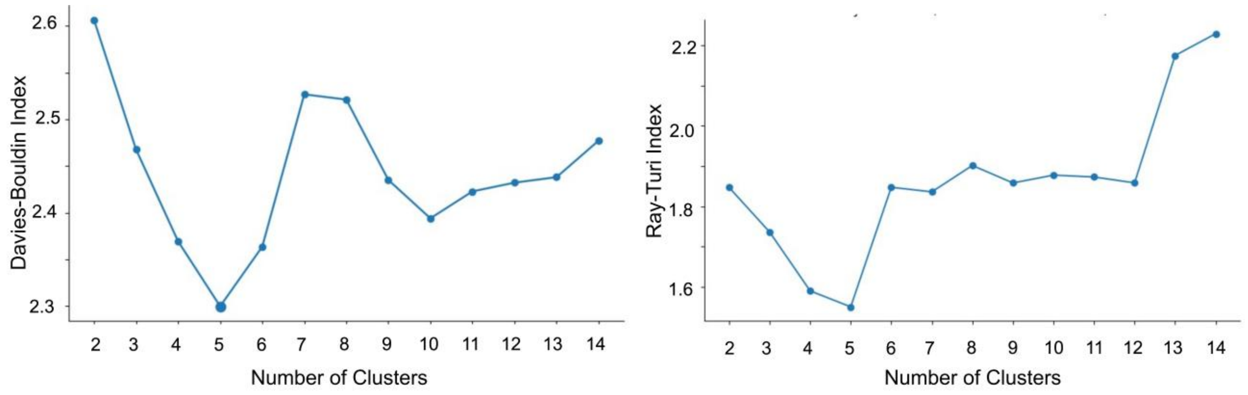


Figure S1. Selection of the optimal number of co-activation patterns. (A) Optimal number calculated by the Davies–Bouldin Index. (B) Optimal number calculated by the Ray–Turi Index.

**Temporal Dynamics of Dwell Time**

To further characterize the temporal evolution of post-learning brain dynamics, we examined the mean dwell time (DW) of the fear-learning–related brain state across the post-conditioning resting period. Using the same sliding-window approach described in the main text, the resting-state time series were segmented into windows of varying lengths, and within each window we computed the average duration for which the fear-learning–related state was continuously maintained.

Across all temporal resolutions, DW exhibited a progressive increase as a function of time following conditioning. Early windows were characterized by shorter dwell durations, whereas later windows showed systematically prolonged state persistence. This temporal profile closely paralleled the time-dependent increase observed for the occurrence rate (OCR) reported in the main results.

Importantly, whereas OCR reflects the frequency with which a state is entered, DW quantifies how long that state is sustained once engaged. The convergence of these two independent dynamic metrics therefore indicates that post-learning neural dynamics are characterized not only by more frequent recruitment of the fear-learning–related brain state, but also by greater temporal stability of that state over time. Together, these supplementary findings provide convergent evidence for a temporally graded strengthening of this state during the post-conditioning resting period.

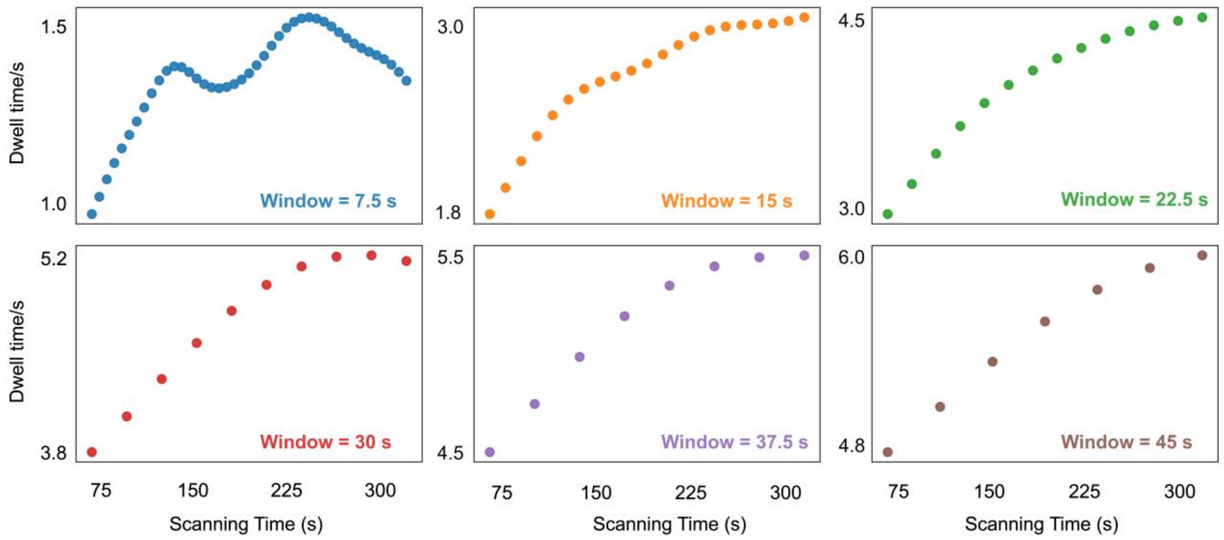


Figure S2. Temporal tracking of fear-learning–induced brain-state engagement during post-conditioning rest.

### Connectivity Reorganization of the Fear-Learning–Related Brain State

To characterize the specific connectivity architecture underlying fear-learning–induced brain reorganization, we quantified state-specific functional connectivity differences between post- and pre-conditioning resting scans for the identified fear-learning–related brain state (State 1). For each participant, connectivity change matrices were computed as the raw difference between post-conditioning and pre-conditioning connectivity (POST − PRE), without additional normalization, and these individual matrices were used as predictors in the main multivariate analyses (Fig. S3A).

To identify connections showing consistent learning-related modulation at the group level, we conducted paired t-tests for each edge across participants. Statistical thresholds were set at three significance levels ($p < 0.05$, $p < 0.01$, and $p < 0.005$, uncorrected) to visualize the spatial pattern of connectivity reorganization across increasingly stringent criteria (Fig. S3 B–D).

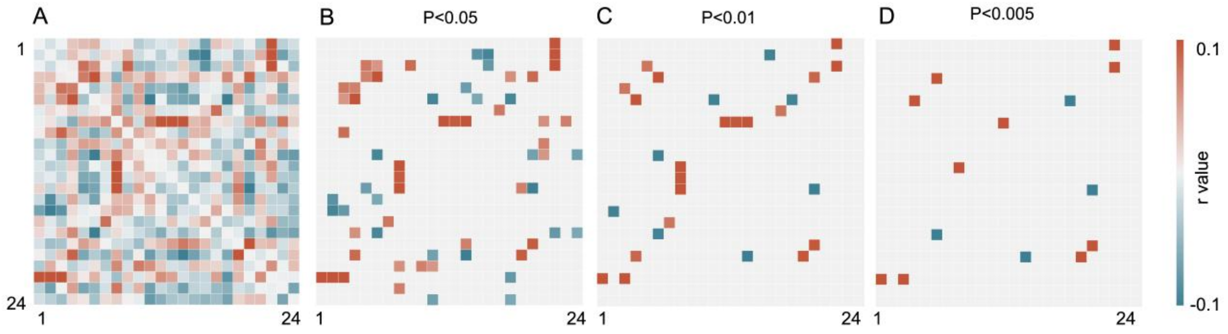


Figure S3. Connectivity reorganization of the fear-learning–related brain state. (A) Group-level visualization of functional connectivity reorganization for the fear-learning–related brain state, computed as the raw difference between post-conditioning and pre-conditioning connectivity matrices (POST − PRE). (A) Group-level visualization of functional connectivity reorganization for the fear-learning–related brain state, computed as the raw difference between post-conditioning and pre-conditioning connectivity matrices (POST − PRE).

### Group-Level Activation Patterns During Extinction Memory Expression After TMS

To characterize the spatial distribution of neural responses during extinction memory expression, we computed group-averaged activation maps across all participants for Day 3 recall and renewal tasks under TMS-modulated extinction. For each condition, activation values were extracted for each of the 24 predefined threat-circuit regions of interest (ROIs) and averaged across participants. ROIs were then ranked according to activation magnitude, and color intensity was assigned proportionally to activation strength to visualize regional contribution profiles.

During both recall and renewal conditions, activation patterns showed a consistent spatial organization. Regions exhibiting the strongest positive activation included visual cortex (VIS), anterior insula subdivisions (dAI and vAI), inferior frontal gyrus opercular part (IFGoper), and dorsal anterior cingulate cortex (dACC), indicating prominent recruitment of sensory-salience–control circuitry during extinction memory expression. In contrast, regions showing relatively reduced activation included medial prefrontal and hippocampal structures, with the ventromedial prefrontal cortex (vmPFC), anterior hippocampus (aHPC), and basolateral amygdala (blAMY) among the most suppressed regions.

Comparing recall and renewal revealed partially overlapping yet distinct activation rankings. While both conditions shared strong engagement of sensory–salience regions, renewal exhibited relatively stronger modulation of hippocampal–limbic and motor-related regions, suggesting differential neural expression profiles across extinction memory retrieval contexts.

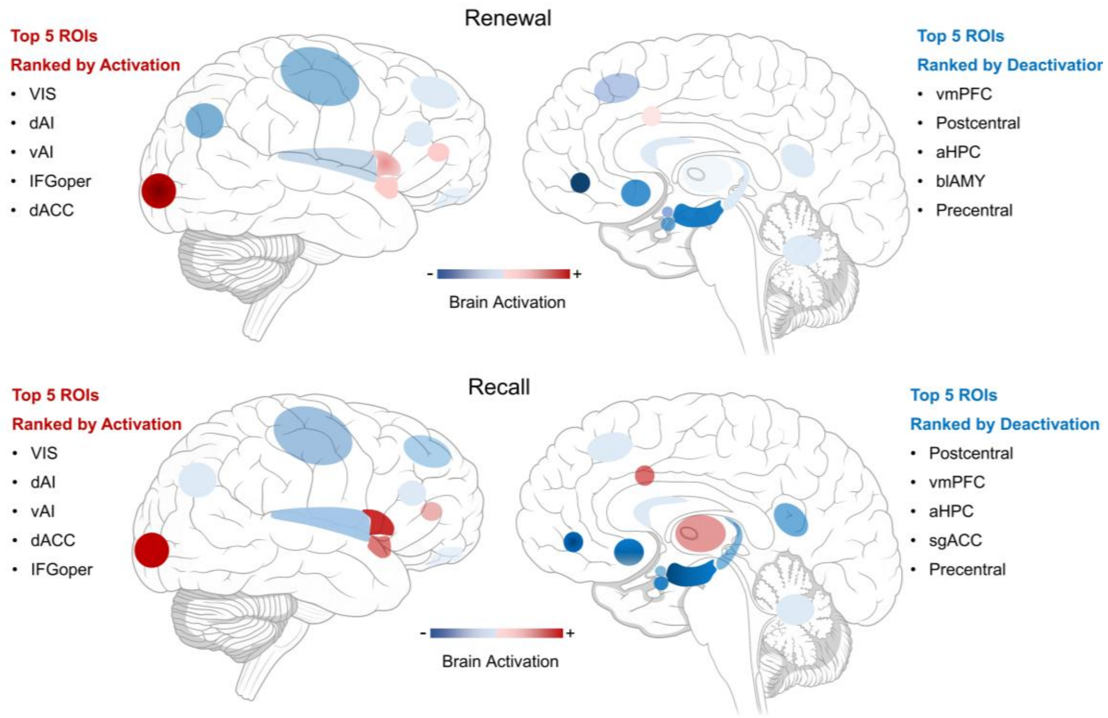


Figure S4. Spatial distribution of mean ROI activation during extinction memory expression following TMS-modulated extinction. Top panel for renewal condition, bottom panel for recall condition.

### Spatial activation profile of the fear-learning–induced brain state

To characterize the spatial configuration of the fear-learning–related brain state identified in the CAP analysis (State 1), we examined its mean activation pattern across the 24 threat-circuit regions. This state exhibited a distributed activation profile spanning medial temporal, insular, cingulate, and sensorimotor regions, indicating coordinated recruitment of multiple functionally distinct components of the threat-processing system.

Ranking regions according to activation magnitude revealed that the strongest contributors were the anterior hippocampus (aHPC), posterior insula (PI), posterior hippocampus (pHPC), precentral gyrus, and basolateral amygdala (blAMY), followed by centromedial amygdala (cmAMY), postcentral gyrus, and angular gyrus. These regions collectively encompass structures implicated in fear memory encoding and contextual representation (hippocampus), threat salience and interoceptive monitoring (insula), affective and associative learning (amygdala), and motor preparation (sensorimotor cortices). The broad spatial distribution indicates that this state reflects a coordinated whole-circuit configuration rather than focal activation within any single region.

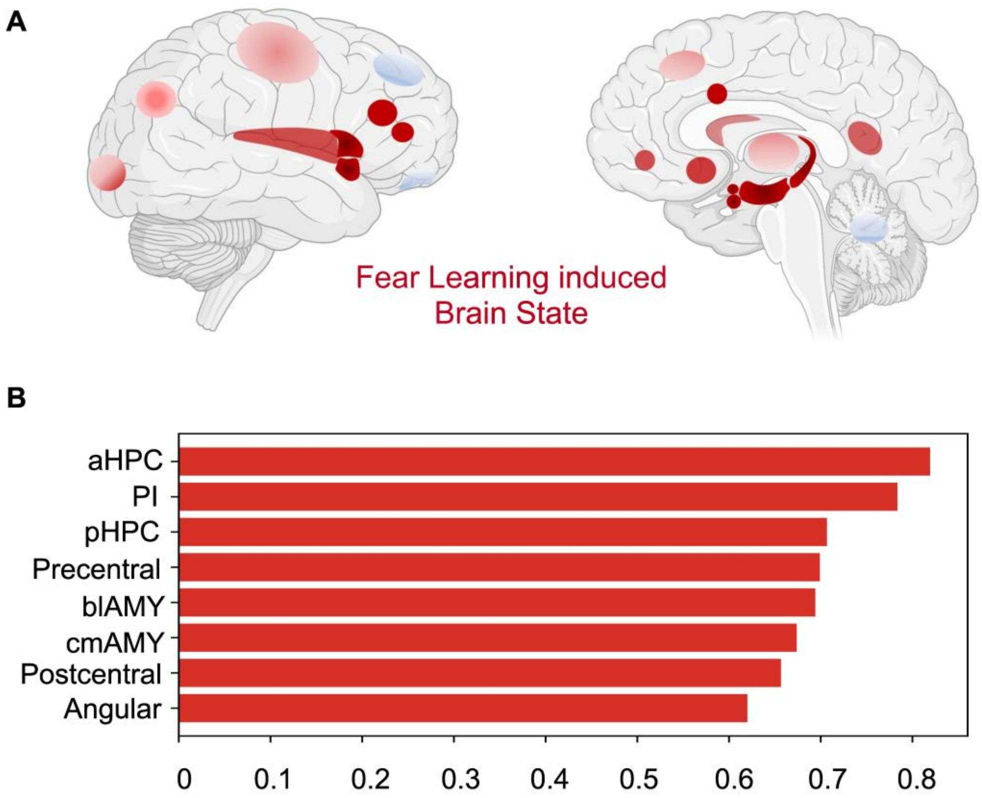


Figure S5. Spatial activation pattern of the fear-learning–induced brain state. (A) Surface and medial views showing group-mean activation weights across the 24 threat-circuit ROIs. (B) Ranking of regional contributions based on activation magnitude.

### Consistency of brain–behavior associations across CS conditions

To verify that the observed brain–behavior associations were not driven by cue-specific effects, we separately examined correlations between fear-learning–induced brain reorganization and behavioral scores across four conditions: recall (CS+TMS, CS+NoTMS) and renewal (CS+TMS, CS+NoTMS). Results show that the two CS+ conditions corresponded to distinct conditioned stimuli presented during Day 1 acquisition (CS+1 and CS+2), allowing us to test whether the identity of the conditioned cue influenced the strength of brain–behavior relationships.

Across all four behavioral measures, the brain reorganization variate exhibited highly consistent and significant positive correlations with subjective fear scores (Figure S6). Specifically, correlations were robust for recall under CS+TMS ($r = 0.70$, $p = 6e\text{-}13$), renewal under CS+TMS ($r = 0.71$, $p = 1e\text{-}13$ recall under CS+NoTMS ($r = 0.72$, $p = 1e\text{-}13$ and renewal under CS+NoTMS ($r = 0.73$, $p = 8e\text{-}14$).

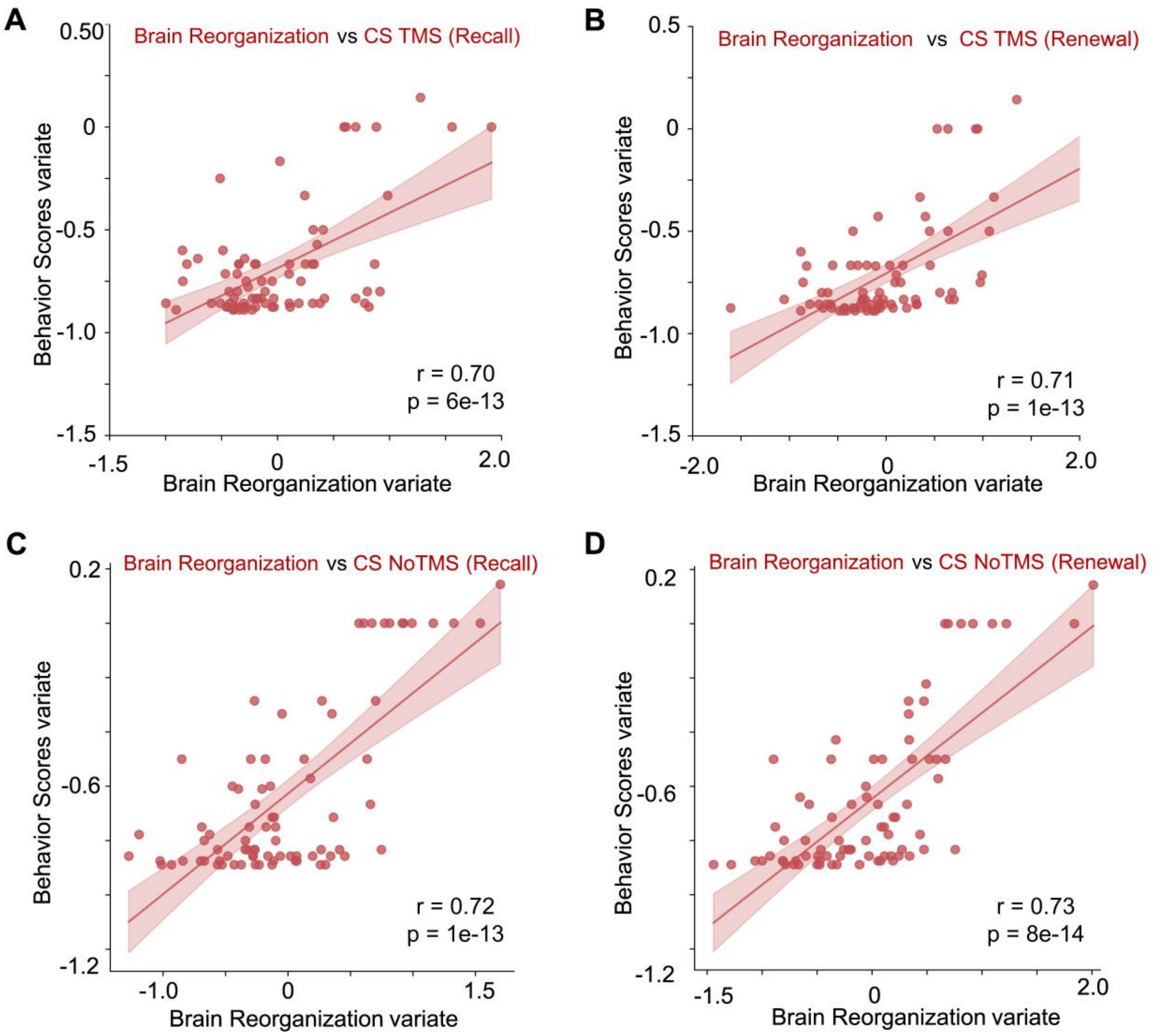

Figure S6. Correlations between fear-learning–induced brain reorganization and behavioral scores across four experimental conditions. (A) Recall under CS+TMS; (B) Renewal under CS+TMS; (C) Recall under CS+NoTMS; (D) Renewal under CS+NoTMS.
